# Age-related differences in white matter: Understanding tensor-based results using fixel-based analysis

**DOI:** 10.1101/751628

**Authors:** Shannon Kelley, John Plass, Andrew R. Bender, Thad A. Polk

**Affiliations:** Department of Psychology, University of Michigan, Ann Arbor, MI 48109, USA; Department of Epidemiology and Biostatistics, Michigan State University, East Lansing, MI 48824, USA

**Keywords:** Aging, diffusion, fixel, tensor, white matter

## Abstract

Aging is associated with widespread alterations in cerebral white matter (WM). Most prior studies of age differences in WM have used diffusion tensor imaging (DTI), but typical DTI metrics (e.g., fractional anisotropy; FA) can reflect multiple neurobiological features, making interpretation challenging. Here, we used fixel-based analysis (FBA) to investigate age-related WM differences observed using DTI in a sample of 45 older and 25 younger healthy adults. Age-related FA differences were widespread but were strongly associated with differences in multifiber complexity (CX), suggesting that they reflected differences in crossing fibers in addition to structural differences in individual fiber segments. FBA also revealed a frontolimbic locus of age-related effects and provided insights into distinct microstructural changes underlying them. Specifically, age differences in fiber density were prominent in fornix, bilateral anterior internal capsule, forceps minor, body of the corpus callosum and corticospinal tract, while age differences in fiber cross section were largest in cingulum bundle and forceps minor. These results provide novel insights into specific structural differences underlying major WM differences associated with aging.

## 1. Introduction

A number of age-related cognitive declines have been associated with changes in cerebral white matter (e.g. DeCarli et al. 1995; Gunning-Dixon and Raz 2000; Turken et al. 2008; Ylikoski et al. 1993), but exactly how white matter changes with age is still unclear. One of the most common approaches to studying age-related changes in white matter is diffusion magnetic resonance imaging (dMRI). By quantifying the diffusion of water molecules in white matter tracts, dMRI can be used to infer the geometry and structural properties of white matter fiber bundles. Because white matter fascicles preferentially restrict diffusion perpendicular to their orientation, the orientation and strength of diffusion in individual voxels can be used to estimate the orientations and microstructural properties (e.g., fiber density) of white matter pathways that traverse them. Using this technique, many previous studies have reported significant differences in the white matter of older vs. younger adults.

One of the most widely used models for characterizing diffusion within individual voxels is the diffusion tensor (DT) model. This approach models diffusion as a three-dimensional zero-mean Gaussian distribution, whose parameters can be used to estimate the average magnitude (mean diffusivity; MD), primary orientation (principal diffusion direction; PDD), and anisotropy (fractional anisotropy; FA) of local diffusion. Of particular interest in white-matter neuroimaging is the FA parameter, which estimates the directional coherence of diffusion within a voxel. FA is sensitive to many influences, including differences in fiber cohesion, diameter and packing density, as well as extent of myelination (Beaulieu 2002; Le Bihan 1995; Pierpaoli and Basser 1996). However, FA’s sensitivity to multiple biological properties makes it challenging to identify which factor or combination of factors is responsible for any observed effects. Furthermore, because the DT model does not distinguish between distinct fiber pathways that can traverse a voxel in different directions, FA can conflate microstructural differences in individual fiber bundles with differences in local multi-fiber geometry in the presence of crossing fibers (Alexander et al. 2002; Tuch et al. 2002; Wedeen et al. 2005). And crossing fibers appear to be the rule rather than the exception: it is estimated that approximately 60-90% of white matter voxels contain multiple directionally distinct fiber populations (Jeurissen et al. 2013). Moreover, the vast majority of findings on age-related WM differences in the past 20 years rely on DT metrics. Therefore, it is possible that such effects may reflect, at least in part, differences in local multi-fiber geometry, rather than microstructural differences more directly related to transmission capacity along particular pathways.

Table 1 provides a summary of a number of influential studies that used DTI to investigate age effects on white matter (a number of other studies have investigated how best to model FA as a function of age, but did not focus on the key differences between younger and older adults (Hasan et al. 2010; Kochunov et al. 2011; Lebel et al., 2012; Westlye et al. 2010). As shown in the main findings column, there is considerable variability in the tracts that have been found to exhibit lower FA in older adults. The genu of the corpus callosum was found to have lower FA in older adults in most studies, while the cingulum, superior longitudinal fasciculus, inferior longitudinal fasciculus, internal and external capsule, fornix, corona radiata, forceps minor, and sagittal striatum were found to have lower FA in older adults in at least two of seven studies. As the FA metric may reflect multiple white matter properties, a key objective of the present study was to investigate select influences underlying age differences in FA using a combination of alternative diffusion measures of microstructural, macrostructural, and multi-fiber geometric properties of human white matter.

**Table 1.** Summary of previous DTI studies of age effects on white matter adapted from Yap et al. (2013). Regions of interest (ROI), Tract-based spatial statistic (TBSS), Tractography (TG), Voxel-based morphometry (VBM).

| References | Subjects | Age range<br>(mean ± SD) | Image<br>Acquisition | Image<br>Analysis | Main Findings<br>(old relative to young) |
| --- | --- | --- | --- | --- | --- |
| Bennett et al.<br>(2010) | 14 younger adults<br>14 older adults | 18-20 (18.9 ± 0.7)<br>63-72 (67.6 ± 3.1) | 3 T<br>35 diffusion<br>directions<br>2.5 mm slice | TBSS | Lower FA: Frontal,<br>posterior pericallosal,<br>superior longitudinal<br>fasciculus, sagittal<br>striatum, genu of the<br>corpus callosum, external<br>capsule, fornix, anterior<br>pericallosal,<br>anterior/superior corona<br>radiata, cerebellum,<br>cerebral peduncle<br>Greater FA: None |
| Burzynska et al.<br>(2010) | 80 younger adults<br>63 older adults | 20-32 (25.7 ± 3.2)<br>60-71 (64.8 ± 2.9) | 1.5 T<br>12 diffusion<br>directions<br>2.5 mm slice | TBSS | Lower FA: anterior,<br>superior and posterior<br>corona radiata, white<br>matter of the superior,<br>inferior, middle, frontal<br>and straight gyri, white<br>matter of the precuneus<br>and superior parietal<br>lobule, cingulum (mainly<br>dorsal), fornix, forceps<br>minor and major, external<br>capsule, internal capsule,<br>sagittal striatum,<br>parahippocampal white<br>matter<br>Greater FA: None |
| Davis et al.<br>(2009) | 20 younger adults<br>20 older adults | n/a (20.04 ± 2.5)<br>n/a (68.89 ± 5.3) | 3 T<br>15 diffusion<br>directions<br>2.0 mm slice | TG<br>TBSS | Lower FA: genu and<br>splenium of the corpus<br>callosum, the cingulum<br>bundle, inferior<br>longitudinal fasciculus,<br>uncinate fasciculus<br>Greater FA: None |
| Giorgio et al.<br>(2010) | 37 younger adults<br>19 mid-adults<br>10 older adults | 23.0-40.2 (n/a)<br>41.0-59.6 (n/a)<br>60.0-81.6 (n/a) | 1.5 T<br>60 diffusion<br>directions<br>2.5 mm slice | TBSS<br>VBM | Lower FA: Anterior<br>thalamic radiations,<br>external capsule, anterior<br>limb of internal capsule,<br>cerebral peduncle and<br>cerebellum corona radiata,<br>superior longitudinal<br>fasciculus, forceps minor,<br>inferior fronto-occipital<br>fasciculus<br>Greater FA: None |
| Hugenschmidt et<br>al. (2008) | 20 younger adults<br>21 mid-adults<br>23 older adults | 18-38 (28.30 ± 6.3)<br>39-64 (47.57 ± 7.6)<br>65-90 (71.17 ± 4.3) | 1.5 T<br>15 diffusion<br>directions<br>3.0 mm slice | ROI<br>VBM | Lower FA: Genu and body<br>of the corpus callosum,<br>forceps minor, superior<br>and inferior frontal gyrus,<br>centrum semiovale, optic radiations and external capsule, corticospinal tract<br>Greater FA: n/a |
| Kennedy and Raz (2010) | 77 adults | 19-84 ($56.49 \pm 16.80$) | 1.5 T<br>6 diffusion directions<br>3 mm slice | ROI | Lower FA with age: Genu and splenium of corpus callosum, internal capsule posterior limb, frontal, parietal, temporal occipital<br>Greater FA with age: None |
| Michielse et al. (2010) | 69 adults | 22-84 ($46.9 \pm 17.8$) | 1.5 T<br>6 diffusion directions<br>1.5 mm slice | TG | Lower FA with age: Genu and ventromedial prefrontal white matter, crus of the fornix, uncinate fasciculus.<br>Stable FA with age: Body of the corpus callosum, rostral, dorsal, and parahippocampal cingulum<br>Greater FA with age: Splenium of corpus callosum (starting from mid-50s onward) |
| Pfefferbaum et al. (2005) | 10 younger adults<br>10 older adults | 22-37 ($28.6 \pm n/a$)<br>65-79 ($72.2 \pm n/a$) | 3 T<br>6 diffusion directions<br>2.5 mm slice | VBM | Lower FA: Superior longitudinal fasciculus, inferior longitudinal fasciculus, anterior cingulate bundle, middle frontal gyrus, frontal forceps and genu<br>Greater FA: n/a |
| Salat et al. (2005) | 15 younger adults<br>9 mid-adults<br>14 older adults | 21-37 ($26.3 \pm n/a$)<br>42-59 ($51.8 \pm n/a$)<br>65-76 ($70.9 \pm n/a$) | 1.5 T<br>6 diffusion directions<br>2 mm slice | ROI | Lower FA: Genu of the corpus callosum, bilateral deep frontal white matter, posterior limb of internal capsule, medial orbitofrontal white matter and posterior periventricular<br>Greater FA: n/a |

Specifically, we used fixel-based analysis (FBA) to assess age-related differences at the level of individual fiber segments (“fixels”) and individual voxels (typically containing multiple fixels). FBA employs a spherical harmonic representation of diffusion that can more readily represent complex multi-fiber geometry than tensor-based models, allowing for anatomically informative metrics to be separately estimated for distinct fiber populations within a voxel (Tournier et al. 2007; Wilkins et al. 2015). Fixel-based analysis utilizes constrained spherical deconvolution (CSD) to estimate fiber orientation distribution functions (fODFs) within each voxel, and then segments fODFs into orientationally distinct lobes corresponding to distinct fiber populations (“fixels”) within a voxel. Figure 1 displays what a tensor model and a fOD model might look like for a single voxel that contains crossing fibers. The fOD model is able to represent multiple orientationally distinct fiber populations, exemplifying its advantage in regions with crossing fibers. This approach allows the diffusion associated with individual fiber segments to be estimated independently by measuring the size of their respective fODF lobes, effectively deconfounding local multi-fiber geometry and tissue microstructure metrics.

**Figure 1.**
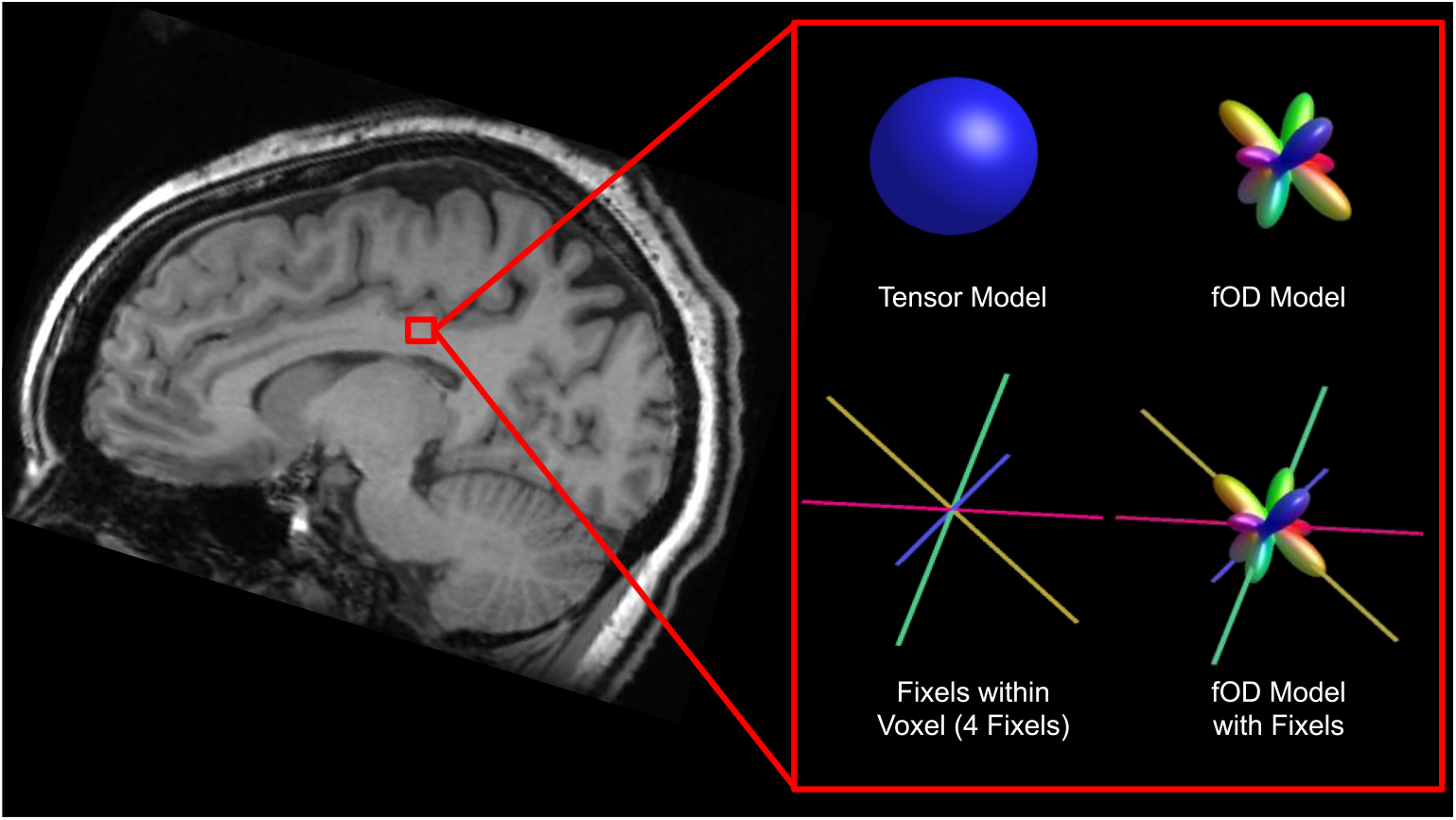
Tensor and fOD model for a single voxel with corresponding fixels, with location displayed on T1 image of corresponding subject.

FBA can be used to estimate both the cross-section of individual fixels (the fiber cross-section or FC) and the density of fibers with a fixel (i.e., fiber density or FD; Raffelt et al. 2017).

Furthermore, the product of these two measures (the fiber density and cross-section or FDC) provides an estimate of the total number of fibers in a fiber bundle. Fixel-based analysis can also be used to estimate the so-called complexity (CX) of the multi-fiber organization within a voxel (Riffert et al. 2014), with complexity being low if a single fiber bundle is present and high in the presence of multiple crossing fibers with similar fiber density.

Figure 2 illustrates the different metrics. The far left of the figure illustrates fODFs of increasing complexity from three different voxels. Complexity measures the relative density of non-primary fixels versus the primary (largest) fixel present in each voxel, ranging from zero (single fixel present) to one (all fixels in the voxel have the same fiber density). The fODF on the top of the figure would be consistent with a single fiber bundle passing through it and therefore has low complexity. Conversely, the fODF on the bottom would be consistent with a voxel containing multiple crossing fibers of similar density and therefore has high complexity.

**Figure 2.**
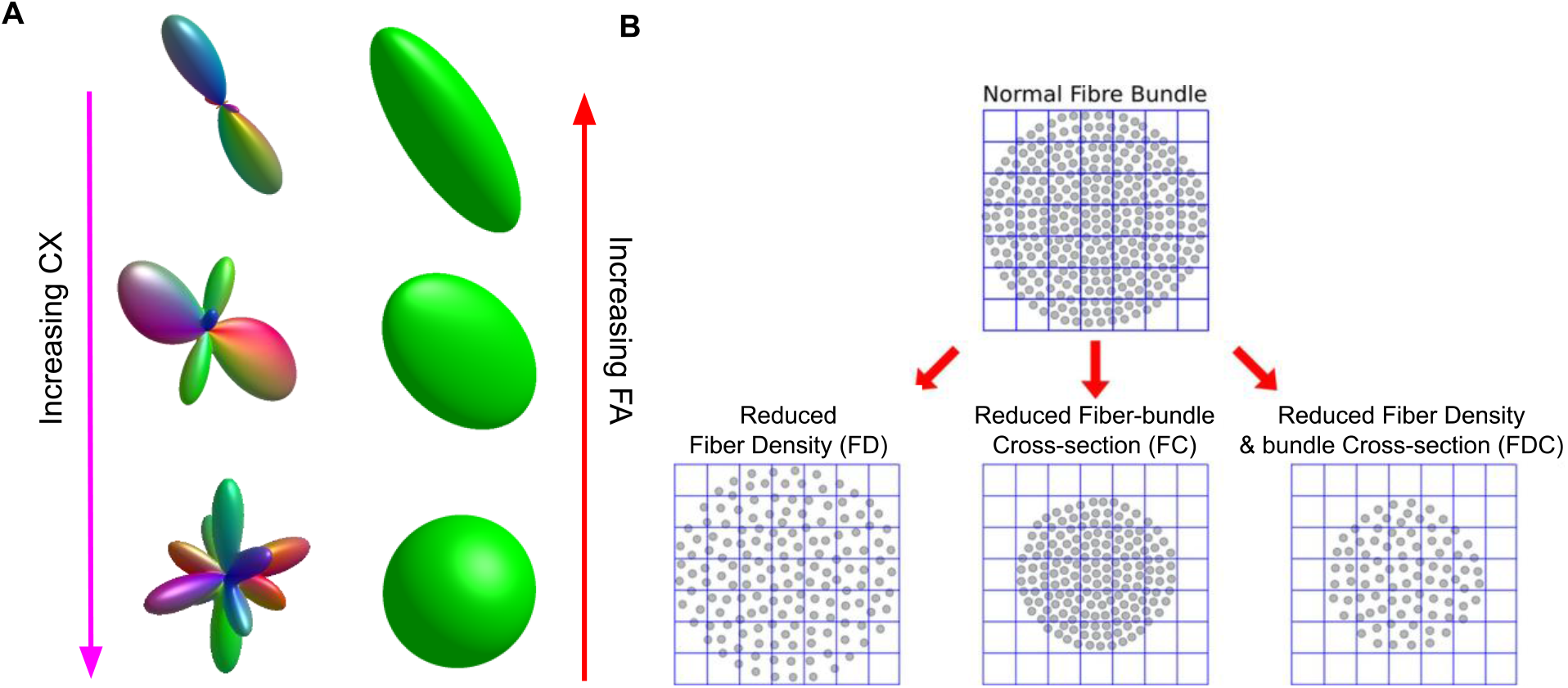
Graphical illustration of complexity (CX), fractional anisotropy (FA), fiber density (FD), fiber crosssection (FC), and fiber density and cross-section (FDC). A) fOD model and tensor model in the same voxels, in which complexity increases from top to bottom while fractional anisotropy decreases. B) An illustration of the FBA metrics FD, FC and FDC adapted from Raffelt et al. (2017).

Immediately to the right of the fODFs in the figure are illustrations of tensor models for the same three voxels. The tensor model at the top is very elongated in the primary direction of water diffusion and very thin in directions perpendicular to that primary direction, leading to a high estimate of fractional anisotropy. Conversely, the tensor model at the bottom is more spherical leading to a low estimate of fractional anisotropy. The fODF for that voxel (on the bottom left of the figure) suggests that there are multiple crossing fibers in that voxel each of which has a distinct direction, but note that because DTI models the data as a single ellipsoid it cannot distinguish the different fiber bundles or estimate their individual properties (e.g., their crosssection or fiber density). The ability to distinguish individual fiber bundles and estimate their properties is one of the main advantages of the fixel-based approach.

On the right of Figure 2 is an illustration (adapted from Raffelt et al. 2017) of the specific metrics computed by the fixel-based analysis: fiber cross-section, fiber density, and the product of cross-section and density. On the top is a cutaway view through a fiber bundle with individual fibers represented by dots. The FBA measure of fiber cross-section (FC) is an estimate of the size of the entire fiber bundle (i.e., the area of the circular area containing individual fibers/dots). The FBA measure of fiber density (FD) is an estimate of the number of individual fibers per unit area (i.e., how densely packed the individual fibers/dots are in the fiber bundle). Finally, the FBA measure of fiber density and cross-section (FDC) is simply the product of the FD and the FC measures. Fiber bundles that are both thick and densely packed will therefore have the highest FDC.

The fiber bundle immediately below and to the left has a lower fiber density (but the same fiber cross-section) as the fiber bundle above it. The fiber bundle directly below and in the middle has a smaller fiber cross-section (but the same fiber density) as the fiber bundle above it. Finally, the fiber bundle below and to the right has a lower fiber density and lower fiber cross section than the fiber bundle above it.

By examining the number and relative size of fixels within each voxel, the fixel-based approach can be used to identify voxels where age effects on FA could potentially be explained by differences in local multi-fiber organization within a voxel, rather than structural differences in individual fiber populations. Because FA is differentially influenced by the strength of diffusion along the principal axis of diffusion versus the two orthogonal axes, decreases in FA can reflect both decreased diffusivity in fiber populations relatively parallel to the principal axis and/or increased diffusivity in relatively “off-axis” populations (Grazioplene et al. 2018). For example, the decreases in FA on the left of Figure 2 appear to be primarily due to this kind of off-axis increase in diffusivity. And to the extent that FA reflects the relative density of on-versus off-axis fiber bundles, age-related changes in FA cannot be unambiguously attributed to changes in a single fiber bundle versus changes in the number of differentially oriented fiber bundles (e.g., crossing fibers).

To assess the extent to which observed age differences in FA could be attributed to changes in such multi-fiber organization, we examined the relationship between FA and the multi-fixel “complexity” (CX) metric. As Figure 2 suggests, increases in complexity might be expected to be associated with declines in FA, and consistent with that intuition previous studies have reported strong negative correlations between FA and CX in a single subject (Riffert et al. 2014), and in a study of schizophrenia (Grazioplene et al. 2018). These results demonstrate that FA (and group differences in FA) can indeed be influenced by the multi-fiber composition of white matter populations within a voxel. We therefore wondered if some of the age-related differences in FA that have been reported in previous studies could be due to age-related differences in crossing fibers and complexity. To our knowledge, that question has not yet been examined.

Here, we used FA, CX, and fixel-level metrics to characterize age-related WM changes throughout the whole brain and in 16 canonical WM pathways. After first testing for age-related differences in FA, we examined correlations between FA and CX in regions exhibiting significant age effects and then tested for age-related effects on FA again after statistically controlling for CX. This allowed us to assess whether observed age effects on FA could plausibly be attributed to changes in local multi-fiber organization. Then, to more directly investigate structural differences between older and younger adults at the level of individual fiber segments, we used FBA to identify age-related differences in estimated fiber density (FD), fiber cross-section (FC), and their product (FDC) in individual fixels throughout the brain (Raffelt et al. 2012).

## 2. Methods

### 2.1 Participants

Participants for this study were recruited as part of the Michigan Neural Distinctiveness (MiND) study (Gagnon et al. 2019). All participants were healthy (no debilitating physical conditions, mental illness, or head trauma), were right-handed and were free of significant cognitive impairment (specifically, they all had a composite overall cognition score > 85 on the NIH Toolbox Cognition Battery; Weintraub et al. 2013). Participants were separated into 25 younger adults (19-29 years old) and 45 older adults (65-87 years old). Participants were excluded if they had motor control or hearing problems, had current depression or anxiety, had a history of drug or alcohol abuse or drank more than 4-6 alcoholic beverages per week (4 for women and 6 for men). Complete information regarding exclusion criteria is reported in Gagnon et al. (2019). All study procedures were approved by the University of Michigan Medical School Institutional Review Board. Participant demographics are displayed in Table 2.

**Table 2.** Age, gender, and education of the study participants.

|  | Younger Adults<br>n = 25 | Older Adults<br>n = 45 | Statistic |
| --- | --- | --- | --- |
| Mean age (SD) | 23.32 (3.06) | 70.69 (4.95) | $t=43.35$<br>$P<.001$ |
| Gender (%) |  |  |  |
| Male | 10 (40) | 17 (37.78) | $\chi^2=.034$ |
| Female | 15 (60) | 28 (62.22) | $P=.855$ |
| Mean years of education (SD) | 15.72 (1.34) | 16.91 (2.10) | $t=2.56$<br>$P=.013$ |

### 2.2 Imaging Acquisition and Processing

MRI data were collected using a 3T General Electric Discovery Magnetic Resonance System with an 8-channel head coil at the University of Michigan’s Functional MRI Laboratory. The dMRI data were collected with a diffusion-weighted 2D dual spin echo pulse sequence with the following parameters: Repetition Time (TR) = 7250 ms; Echo Time (TE) = 2.5 ms; Field of View (FOV) = 240 x 240 mm; 32 diffusion directions; 60 axial slices with thickness = 2.5 mm (0.9375 mm in-plane resolution) and 0.1 mm spacing. Five volumes without diffusion weighting (b = 0 s/mm^2^) and 32 diffusion-weighted volumes (b = 1000 s/mm^2^) were collected. Acquisition time was approximately 10 minutes. Two scans were collected with the previously described properties with opposite phase-encoding. T1-weighted structural images were collected with the following parameters: TR = 3173.1 ms; TE = 24.0 ms; Inversion Time (TI) = 896 ms; flip angle = 111°; FOV = 220 x 220 mm; 43 axial slices with thickness = 3 mm and no spacing; acquisition time = 100 seconds.

Diffusion Magnetic Resonance Image (dMRI) processing followed the published steps outlined in the MRtrix3 user manual (https://mrtrix.readthedocs.io/en/3.0_rc2/fixel_based_analysis/ss_fibre_density_cross-section.html). Details of our pipeline can be found on GitHub (https://github.com/umich-tpolk-lab/fba_paper). This included preprocessing the data to correct for susceptibility distortion and motion and to apply an eddy current correction, all using FSL’s eddy_correct/TOPUP tools (Jenkinson et al. 2012). Each scan was individually examined for major artifacts. Bias field correction was performed using ANTS N4 (http://picsl.upenn.edu/software/ants/). dMRIs were intensity normalized across subjects based on the median b = 0 s/mm^2^ intensity within a white matter mask and resampled to an isotropic voxel size of 1.3 mm using b-spline interpolation. A bias field correction was also applied (Tournier et al. 2019).

Constrained spherical deconvolution (CSD; “Tournier” algorithm) was used to compute fiber orientation distributions (FODs) (Tournier et al. 2007). A group average response was used to estimate FODs in all subjects, as described in Raffelt et al. (2012). A white matter template fixel mask was generated with a peak amplitude threshold of 0.15. Whole brain probabilistic tractography was then performed on the FOD template generating 20 million streamlines and spherical-deconvolution informed filtering of tractograms (SIFT) was applied with an output of 2 million streamlines (Smith et al. 2013). SIFT removes individual streamlines so that the density of reconstructed connections is proportional to the fiber density as estimated by the diffusion model, providing a biologically relevant estimation of the density of white matter axons connecting two regions. The remaining streamlines were used to estimate the fixel-fixel connectivity matrix, which was used to inform threshold-free cluster enhancement in the fixel-based analysis (below).

A five-tissue-type (5TT) image was generated from the T1-weighted structural image using FSL. Volumes from the 5TT image were used to estimate regions containing CSF and were included as masks in all analyses to minimize partial voluming effects.

### 2.3 Whole-brain Voxel-Based and Fixel-Based Analysis

The general workflow for processing and analyzing the data is shown in Figure 3. Both the whole-brain DTI analysis and the FBA analysis were performed using MRtrix3. Diffusion tensor images were generated from group-registered diffusion weighted images using an iteratively reweighted linear least squares estimator. Fractional anisotropy (FA) maps were then created for each subject and smoothed using a Gaussian kernel with a standard deviation of 1.3 mm using the mrfilter command. Whole-brain voxel-based analysis was then performed to identify voxels in which FA was significantly different in the young vs. the older participants (with threshold-free cluster enhancement, default parameters: dh=0.1, e=0.5, h=2) using the mrclusterstats command. To control family-wise error (FWE) rate, 5,000 permutations were used to derive a null distribution of maximum cluster-enhanced test statistics across voxels and then the true test statistic for each voxel was compared against that null distribution to determine if it was beyond the 95th percentile of the distribution. Each voxel was then assigned a FWE-corrected *P*-value (Smith and Nichols 2009; Holmes et al. 1996). We localized the effects based on the Catani and Schotten (2012) and Wakana et al. (2007) white matter atlases.

**Figure 3.**
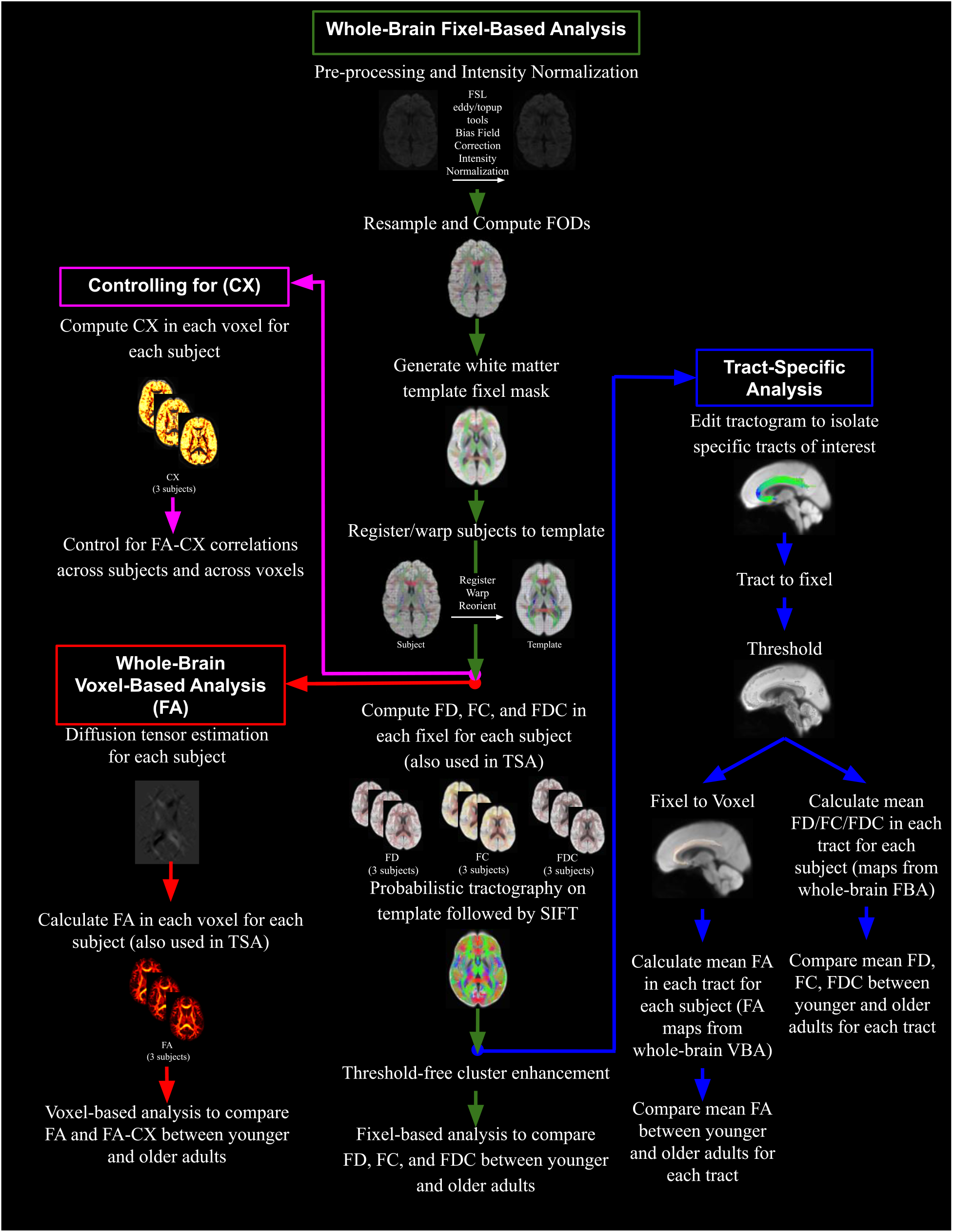
Diagram displaying the workflow for four of the different analyses conducted in the current study.

We also performed a similar whole-brain analysis with MRtrix3 but using fixel-based methods. Fiber density (FD), fiber cross section (FC), and fiber density and cross section (FDC) were calculated for every fixel for each subject. T-tests statistically compared younger to older adults at each white matter fixel. Connectivity-based fixel enhancement (CFE) was used to perform smoothing and cluster-based statistical inference. CFE identifies structurally connected fixels that likely share underlying anatomy using probabilistic tractography. Prior to statistical analysis, tract-specific smoothing was performed, followed by enhancement of the statistical map using threshold-free cluster enhancement based on estimated fixel-fixel connectivity within the population template (Raffelt et al. 2015). CFE was performed using 2 million streamlines and default parameters (smoothing = 10 mm FWHM, C = 0.5, E = 2, H = 3; taken from Raffelt et al. 2015). 5,000 permutations were used to derive a null distribution of maximum cluster-enhanced test statistics across fixels and then the true test statistic for each fixel was compared against that null distribution to determine if it was beyond the 95th percentile of the distribution. Each fixel was then assigned a FWE-corrected *P*-value. We localized the effects based on the Catani and Schotten (2012) and Wakana et al. (2007) white matter atlases.

We used MRtrix’s fixel2voxel command to estimate complexity (CX) at every voxel for each participant. To determine the correlation between FA and CX in voxels with significantly different FA between age groups, we first used the mrdump command to compute FA and CX values for each subject in every voxel that exhibited a significant age difference in FA. We then used an in-house script implemented in MATLAB (R2017b) to determine the correlation between FA and CX values in those voxels. We first computed the average FA across the whole mask in each subject and the average CX across the whole mask in each subject. We then computed the correlation between these measures of average FA and average CX across subjects. Additionally, we calculated the correlation between FA and CX across all voxels in the significance mask within each subject. We then examined the distribution of these correlation coefficients across subjects.

Finally, to estimate age-related effects on FA while controlling for CX, we controlled for complexity differences both across individuals and across voxels within each individual. First, to control for CX across individuals, we regressed mean FA onto mean CX and subtracted each subject’s predicted mean from all their voxels. Next, to control for CX across voxels, we regressed CX onto FA across voxels within each subject’s analysis mask. We then subtracted the slope term of these regressions (CX times the change in FA per unit change in CX) from each subject’s mean-corrected FA image, resulting in FA images with means corrected according to across-subject correlations, and voxel-wise deviations from the mean corrected according to within-subject correlations. This approach ensured that corrections at the within-subject level did not affect subjects’ mean FA, so that that mean differences remaining after between-subjects correction were retained (see Supplemental Figure 1).

### 2.4 Tract-Specific Analyses

We also performed tract-specific analyses using both DTI-based and FBA-based methods. We selected sixteen major tracts for analysis, including canonical pathways included in most white matter parcellations and previous studies of age group differences in the DTI literature. White matter atlases from Wakana et al. (2007) and Catani and Schotten (2012) were used as anatomical guidelines to manually select regions for inclusion and exclusion. The white matter template computed by MRtrix was used for ROI placement so that all subjects’ data were coregistered beforehand. We used the tckedit command in MRtrix3 to extract tracts of interest from the whole brain tractogram and mapped them to fixels using the tck2fixel command (45 degree angular threshold). Tracts were thresholded to only include fixels that had at least 1% of each tract’s streamlines associated with them. The thresholded fixel maps were used for calculating mean FD, FC, and FDC for each tract for each subject. Using the fixel2voxel command in MRtrix, we then converted the thresholded fixel maps to voxel maps. The voxel maps were used to calculate mean FA for each tract for each subject. The following tracts were included: cingulum (separated into retrosplenial, subgenual cingulum and parahippocampal cingulum as described in Jones et al. (2013); the parahippocampal cingulum was further separated into parietal and temporal components), corticospinal tract (separated into superior and inferior), forceps major, forceps minor, fornix, inferior fronto-occipital fasciculus (IFOF), inferior longitudinal fasciculus (ILF), internal capsule, superior longitudinal fasciculus (SLF, separated into SLF I, II and III as described in Schotten et al. 2011) and uncinate fasciculus. SLF I projects to the parietal precuneus and supplementary motor area, SLF II projects to the posterior region of the inferior parietal lobule and the lateral aspect of the superior and middle frontal gyrus, and SLF III projects to the inferior parietal lobule and the posterior region of the inferior frontal gyrus (Catani and Schotten 2012). The parahippocampal cingulum and corticospinal tract were each divided into two components to facilitate comparison with our whole-brain analyses, which indicated distinct local effects along these pathways. Mean FA, FD, FC and FDC were calculated for each tract bilaterally. 10,000 permutations were used to derive a null distribution of maximum *t*-statistics across tests and then the true *t*-statistic for each test was compared against that null distribution to determine if it was beyond the 95th percentile of the distribution.

### 2.5 Visualizing Results

Visualizations of the results were generated using the mrview tool from MRtrix3. The two million streamlines generated from SIFT were cropped to only include fixels or voxels in which the effect of age group was significant as defined by a family-wise error-corrected *P*-value below 0.05. The significant streamline segments are displayed on the white matter population template.

Results for tract specific analysis are also displayed on the white matter population template. For tracts that were found to be significantly different between older and younger adults following a two-sample t-test, Cohen’s d was calculated. The tracts were generated in template space, and are color coded by the calculated Cohen’s d.

## 3. Results

### 3.1 Whole Brain DTI Analysis

Figure 4 displays white matter voxels that exhibited a significantly lower FA value in older adults compared to younger adults. The voxels are colored according to effect size. Older adults exhibited widespread patterns of lower FA values, relative to younger participants, in a variety of white matter tracts including forceps minor, fornix, bilateral IFOF, bilateral SLF I and III, as well as bilateral anterior internal and external capsule. The strongest age effects (i.e., FA in younger > older) were apparent in the fornix and in the genu of the corpus callosum (Figure 4).

**Figure 4.**
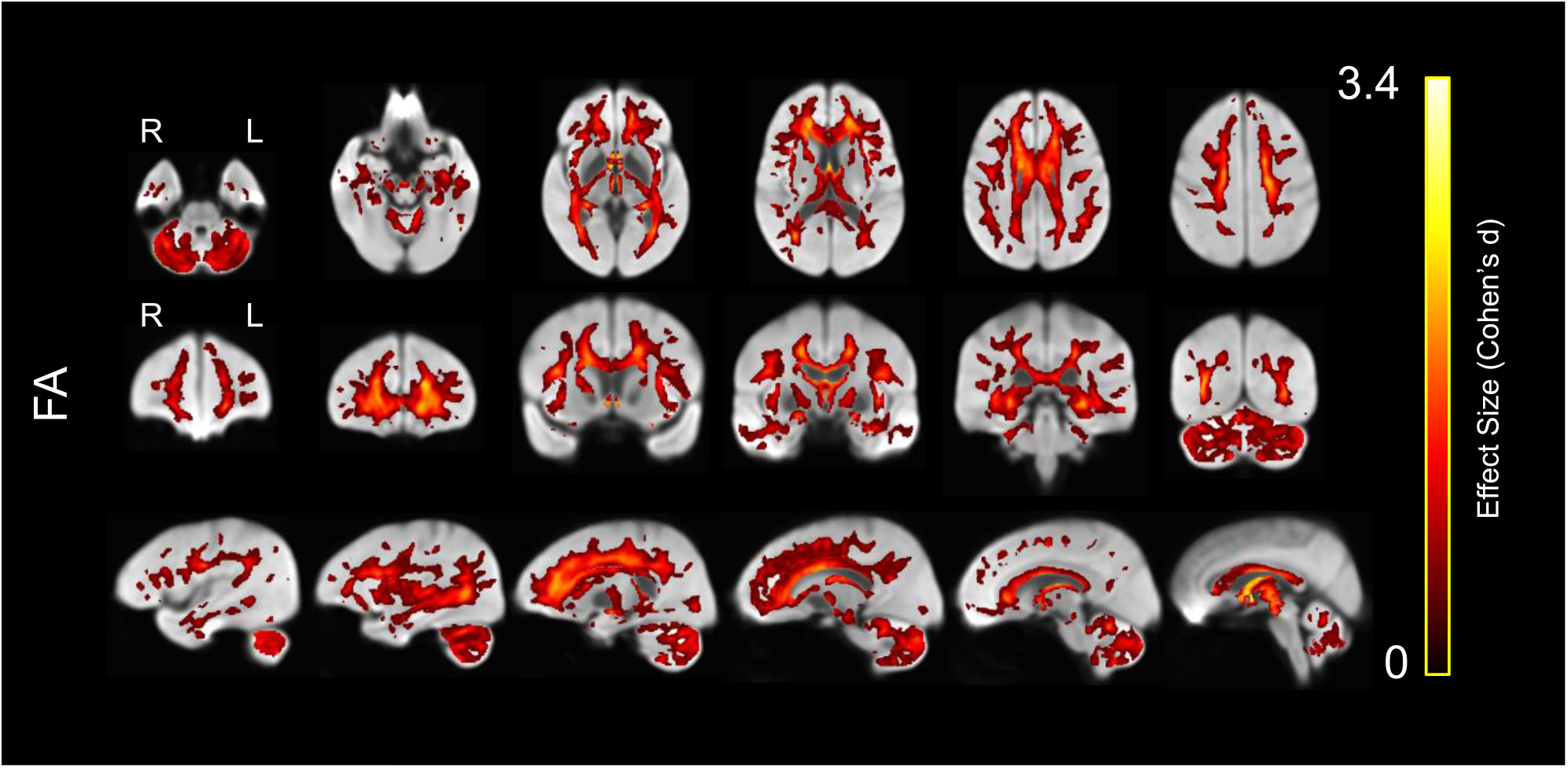
Lower white matter fractional anisotropy (FA) in older compared to younger adults, displayed on the white matter population template. Colors represent voxels in which FA was significantly lower in the older compared with the younger group, with brighter colors representing larger effect sizes (Cohen’s d).

We also evaluated the opposite contrast by evaluating voxels in which older adults had greater FA than younger participants (Figure 5). The voxels in Figure 5 are also colored by the effect size. The regions in which FA was greater for older adults were primarily limited to the superior cerebellar peduncles, external capsule and cingulum bundle.

**Figure 5.**
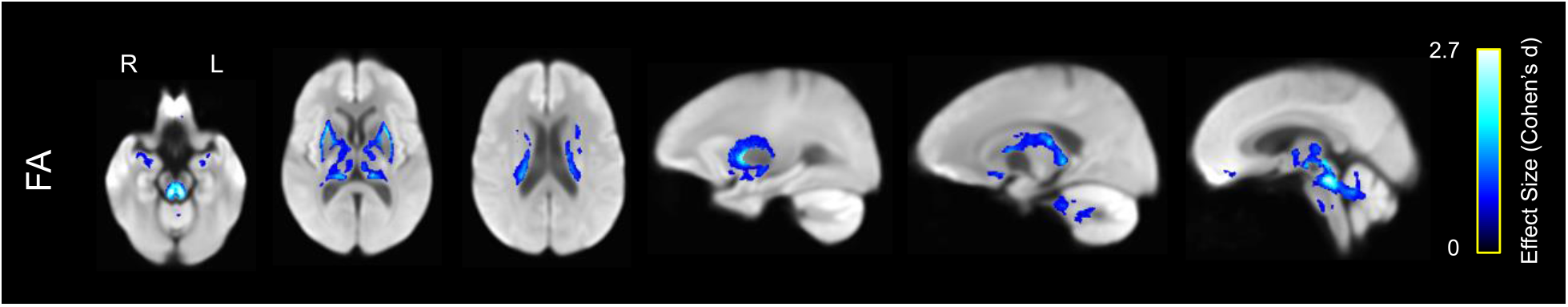
Greater white matter fractional anisotropy (FA) in older compared to younger adults, displayed on the white matter population template. Colors represent voxels in which FA was significantly greater in the older compared with the younger group, with brighter colors representing larger effect size (Cohen’s d).

### 3.2 Tract-Specific DTI Analysis

In addition to whole brain analyses, we also performed tract-specific analyses on 16 tracts (Figure 6). We chose these specific white matter tracts to reflect canonical pathways included in most white matter parcellations and previous studies in the DTI literature comparing age groups.

**Figure 6.**
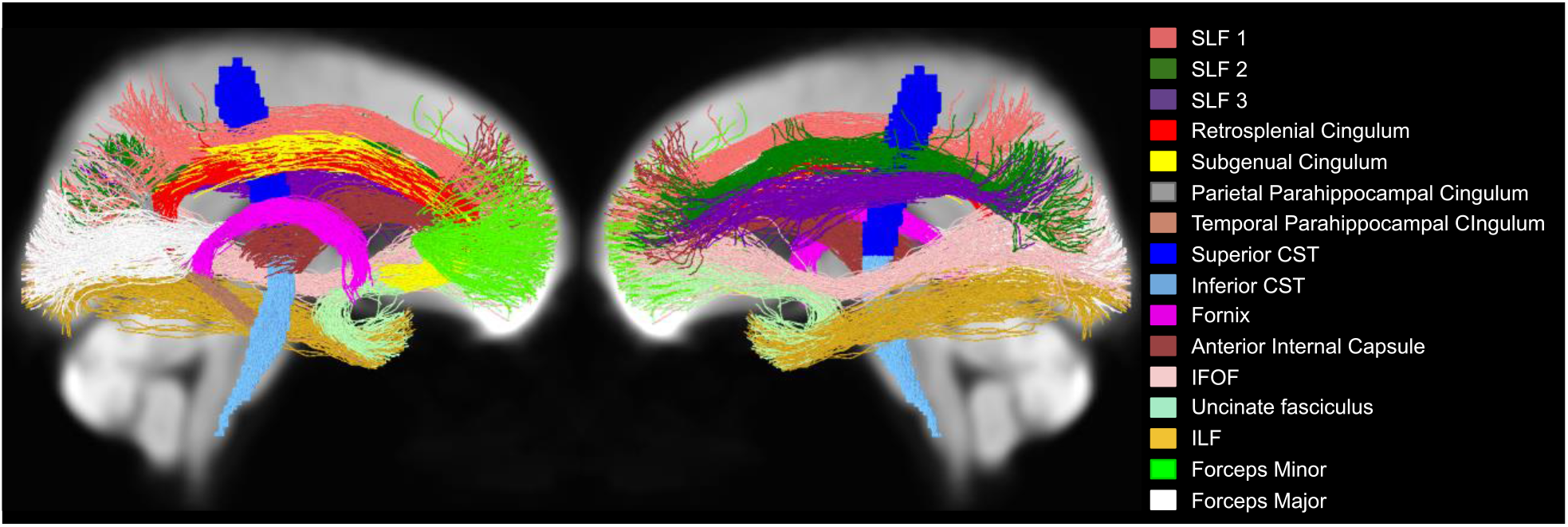
The 16 tracts that were included in the tract-specific analyses, displayed on the white matter population template.

In addition, the parahippocampal cingulum and corticospinal tracts were further divided based on prior work (Jones et al. 2013) and on the results from our whole-brain analysis, which suggested the presence of differences along these pathways.

The age group comparisons of mean FA across tracts showed widespread differences in which older adults mean tract FA was significantly lower than younger adults (Figure 7 and Table 3). Similar to the whole-brain analysis, FA was significantly lower in older vs. younger adults in forceps minor, fornix, bilateral IFOF, bilateral internal capsule, and bilateral SLF I and SLF III (for all, *P*<.001). FA was also lower in SLF II (*P*<.001 for left and *P*=.002 for right), bilateral ILF (*P*=.008 for left and *P*=.002 for right), bilateral parietal parahippocampal cingulum (*P*=.003 for left and *P*=.006 for right) and right uncinate fasciculus (*P*=.022). Mean FA was greater in older adults compared to younger adults in right subgenual (*P*=.002) and right retrosplenial cingulum (*P*=.019) (not shown).

**Table 3.** Results from tract-specific analyses showing significant age differences in FA, CX-corrected FA, FD, FC and FDC with corresponding significant p-values. Analyses in which younger participants exhibited significantly higher values than older participants are colored red and those in which older participants exhibited significantly higher values are colored blue. All analyses used 10,000 permutations to correct for multiple comparisons. Dashes represent no significant difference.

| Tract | FA | FA-CX<br>Correction<br>(Across<br>Subjects) | FA-CX<br>Correction<br>(Across<br>Subjects &<br>Voxels) | FD | FC | FDC |
| --- | --- | --- | --- | --- | --- | --- |
| Left Cingulum retrosplenial | - | - | 0.002 | - | <0.001 | <0.001 |
| Right Cingulum retrosplenial | 0.019 | 0.002 | 0.003 | - | <0.001 | <0.001 |
| Left Cingulum subgenual | - | - | - | - | <0.001 | <0.001 |
| Right Cingulum subgenual | 0.002 | <0.001 | 0.006 | - | <0.001 | <0.001 |
| Left Cingulum parahippocampal parietal | 0.003 | 0.01 | - | - | - | - |
| Right Cingulum parahippocampal parietal | 0.006 | 0.020 | - | 0.023 | - | - |
| Left Cingulum parahippocampal temporal | - | - | - | <0.001 | - | <0.001 |
| Right Cingulum parahippocampal temporal | - | - | - | <0.001 | - | <0.001 |
| Left Corticospinal Inferior | - | - | - | <0.001 | 0.003 | <0.001 |
| Right Corticospinal Inferior | - | - | - | <0.001 | 0.002 | <0.001 |
| Left Corticospinal Superior | - | - | - | - | - | - |
| Right Corticospinal Superior | - | - | - | - | - | - |
| Forceps Major | - | - | - | - | - | - |
| Forceps Minor | <0.001 | <0.001 | <0.001 | <0.001 | - | <0.001 |
| Fornix | <0.001 | <0.001 | <0.001 | <0.001 | <0.001 | <0.001 |
| Left IFOF | <0.001 | <0.001 | - | <0.001 | - | - |
| Right IFOF | <0.001 | <0.001 | - | <0.001 | - | 0.009 |
| Left ILF | 0.008 | - | 0.040 | - | - | - |
| Right ILF | 0.002 | - | 0.009 | - | - | 0.030 |
| Left Anterior Internal Capsule | <0.001 | 0.003 | - | <0.001 | - | <0.001 |
| Right Anterior Internal Capsule | <0.001 | <0.001 | <0.001 | <0.001 | - | <0.001 |
| Left SLF I | <0.001 | <0.001 | - | 0.013 | 0.005 | - |
| Right SLF I | <0.001 | <0.001 | - | - | 0.001 | - |
| Left SLF II | <0.001 | <0.001 | 0.018 | - | - | - |
| Right SLF II | 0.002 | 0.022 | - | - | - | - |
| Left SLF III | <0.001 | 0.004 | <0.001 | - | - | - |
| Right SLF III | <0.001 | <0.001 | <0.001 | - | - | - |
| Left Uncinate Fasciculus | - | - | - | <0.001 | - | <0.001 |
| Right Uncinate Fasciculus | 0.022 | - | - | - | - | 0.030 |

**Figure 7.**
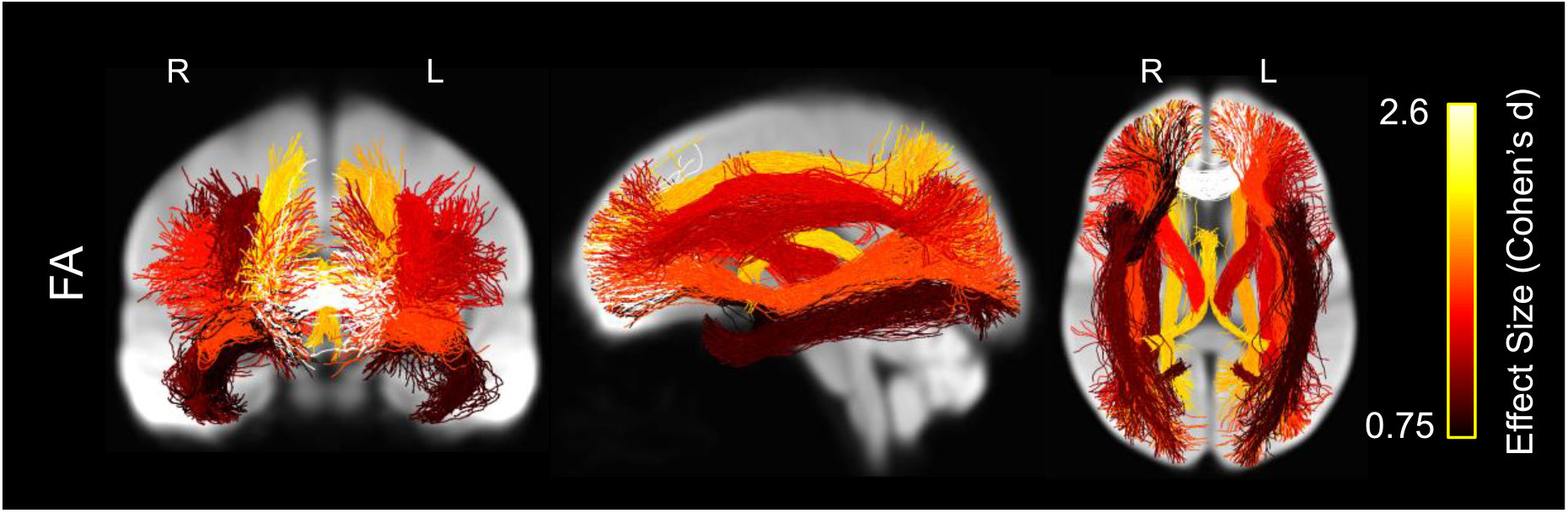
Whole brain projections onto 2-D slices showing white matter tracts in which average fractional anisotropy (FA) was significantly lower in the older vs. younger participants, displayed on the white matter population template. Streamlines within each tract are colored by the tract’s effect size (Cohen’s d), with brighter colors representing greater effect size.

### 3.3 Correlation Between FA and CX

We evaluated the correlation between mean FA and CX within the FA significance mask across subjects (Figure 8). We observed a very strong negative correlation between FA and CX *(r=-* 0.81, *P*< .001). Each participant also exhibited a significant (*P*<.001) negative correlation (mean *r*=−0.72) between FA and CX across voxels in the mask (Figure 8, inset).

**Figure 8.**
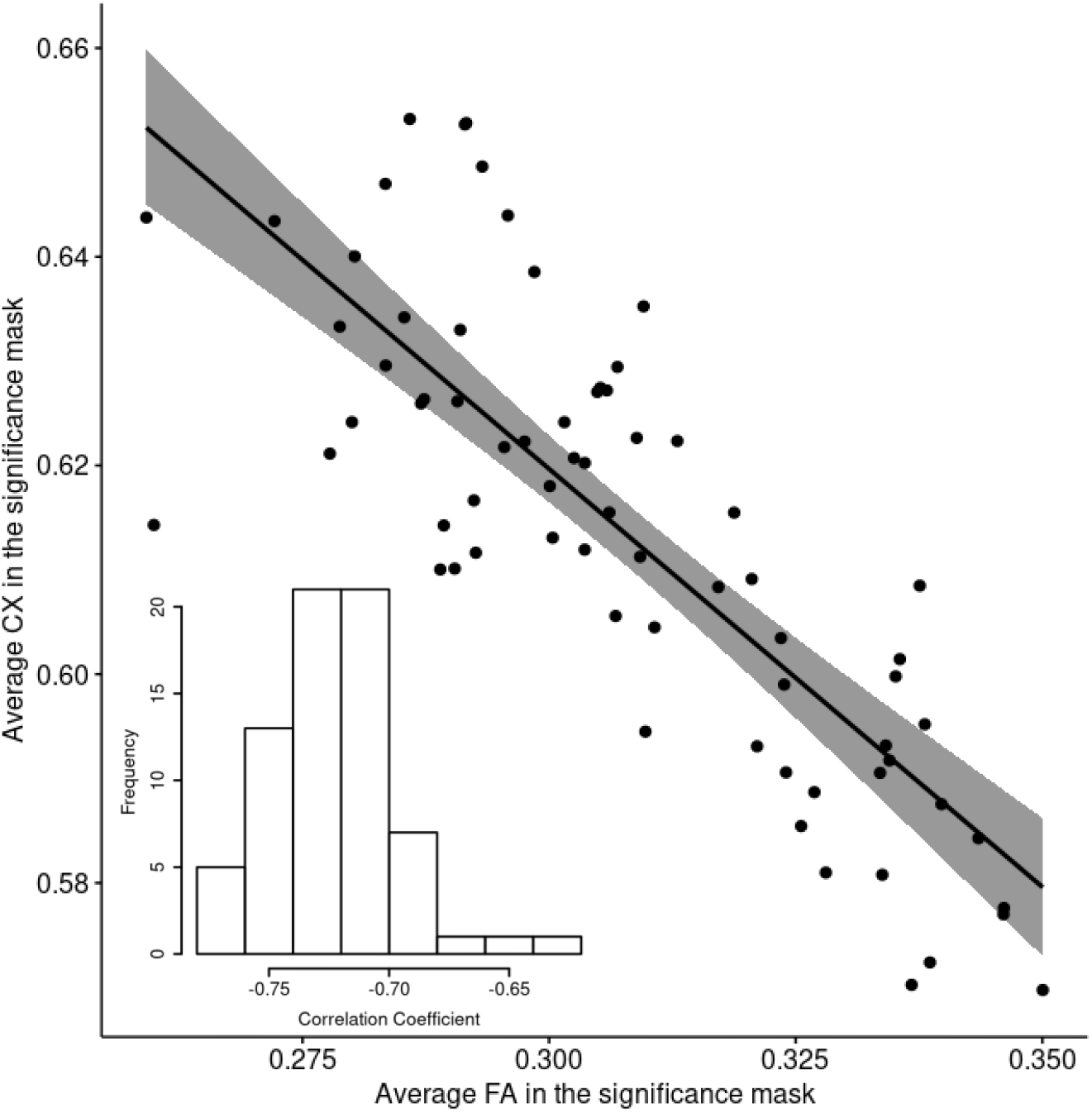
Correlation between average FA and average CX within the significance mask across subjects (*P* < 0.001 and *r* = −0.81). The bottom left corner contains a histogram displaying the frequency of correlation coefficients between FA and CX across voxels within each subject. The mean correlation coefficient was *r* =-0.72.

To further investigate the correlation between FA and CX within voxels, we computed the correlation coefficient at every voxel of the white matter mask across participants (Figure 9). We observed a very strong widespread negative correlation between FA and CX, with minimal areas of no significance or positive correlation. Similar results were obtained in both the young and the older group when they were analyzed separately (supplementary Figures S2 and S3).

**Figure 9.**
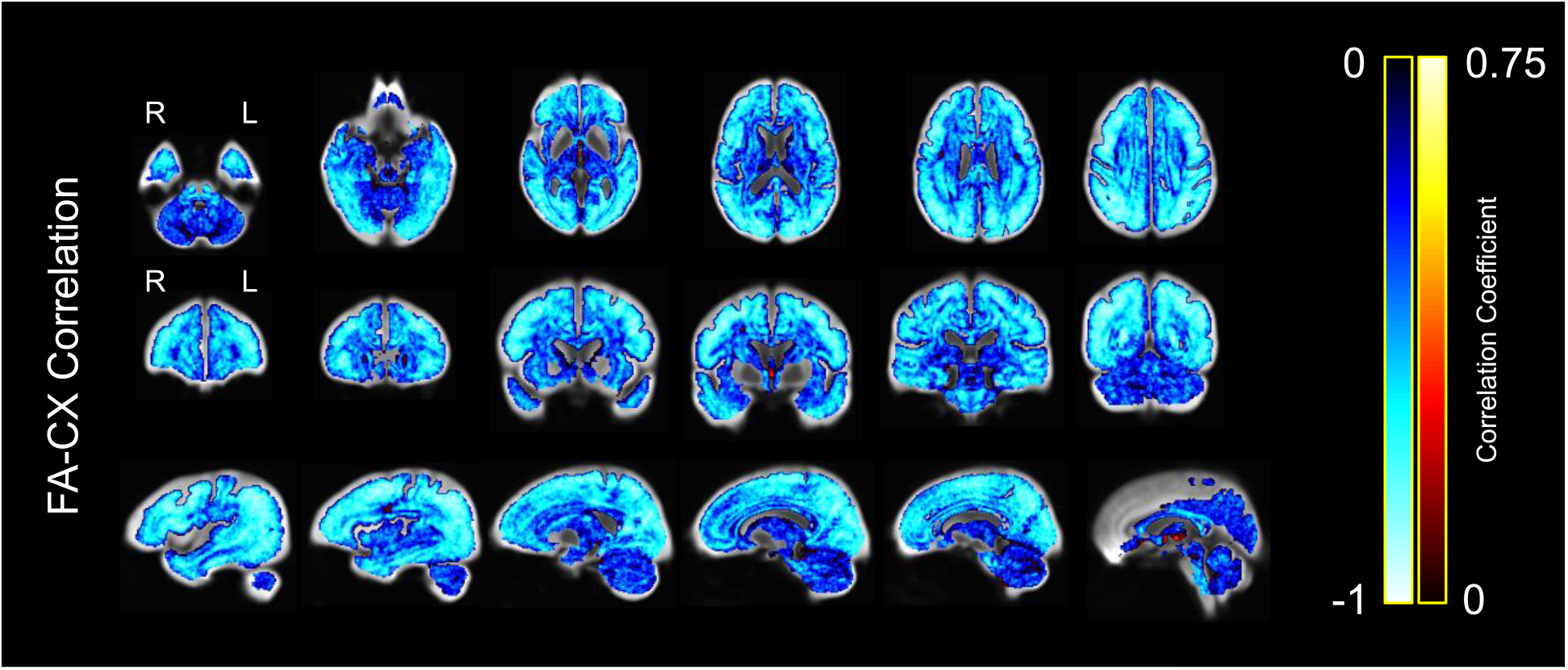
Correlation between FA and CX across subjects at each voxel, displayed on the white matter population template. Colors represent the strength of the negative or positive correlation coefficient for each voxel, with brighter colors representing stronger correlation coefficients.

### 3.4 Whole Brain DTI analysis controlling for CX

To further assess the relationship between CX, FA, and age, we examined CX-FA correlations both across subjects (i.e., correlations between mean CX and FA) and within subjects (i.e., correlations across voxels). Then, we statistically controlled for these correlations to assess how they each contributed to apparent age effects on FA.

Figure 10A shows the mean CX and FA for each individual (points) and the estimated slopes of corresponding within-subject regressions across voxels (lines). Individual points/lines are colored according to age group (blue = young; orange = old). As can be appreciated from the figure, CX and FA were strongly correlated both across subjects (r=-0.54, P<.001) and within subjects (All r<-0.48, all p<.001). Moreover, mean FA and mean CX differed in opposite directions with age (Figure 10A, colored histograms; FA: t[68]=-4.57, *P*<.001; CX: t[68]=3.26, *P*=.002), suggesting that CX-FA correlations likely influenced apparent age effects on FA.

**Figure 10.**
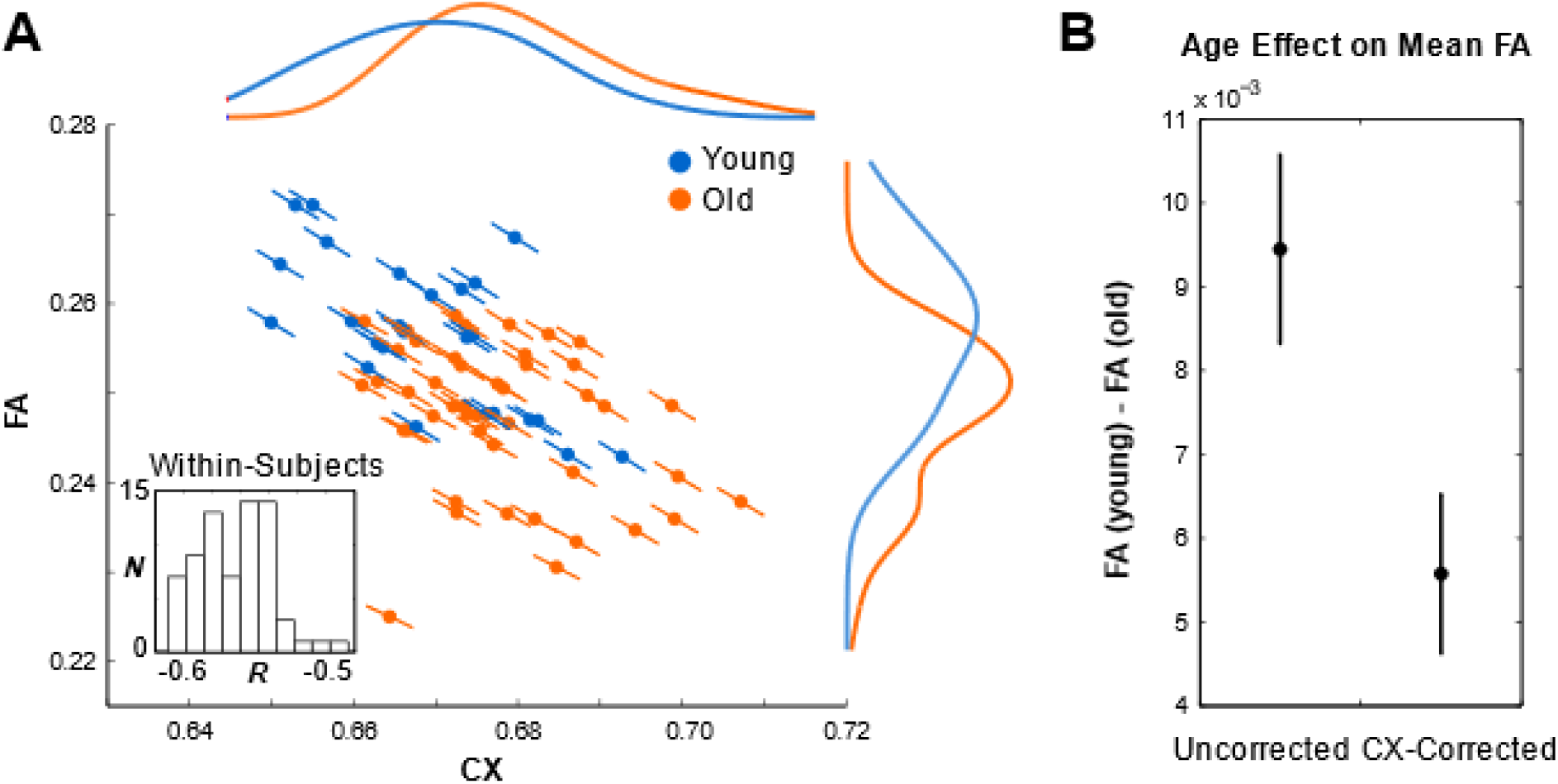
Relationship between CX, FA, and age. (A) Correlations between CX and FA across subject means (colored points) and across voxels within each subject (slopes: colored lines; the inset displays a histogram of the distribution of these within-participant correlation coefficients). Colors indicate age group (blue = young; orange = old). Colored curves show interpolated histograms (kernel density plots) of subject means for each group. (B) Observed age differences in mean FA before (left) and after (right) statistically controlling for CX-FA correlations across subjects.

To more directly examine the influence of CX-FA correlations on observed age differences, we re-examined age effects on FA after statistically controlling for CX-FA correlations, both across subjects and across voxels within subjects. Controlling for mean CX across subjects substantially reduced but did not fully abolish observed age differences in mean FA. Specifically, age differences in mean FA were still significant after controlling for CX (t[68]=2.98, *P*=.004), but the size of the effect was significantly reduced (t[68]=3.26, *P*=.002). As expected, this resulted in a significant reduction in the apparent age effect on FA throughout the entire analysis mask (Fig 11, right, top row), which reduced the spatial extent of significant voxels (Fig 11, left, center row). Whereas the original DTI analysis produced 100,991 significant voxels (25.8% of the analysis mask), controlling for CX-FA correlations across subjects reduced this number to 63,205 voxels (16.1%).

**Figure 11:**
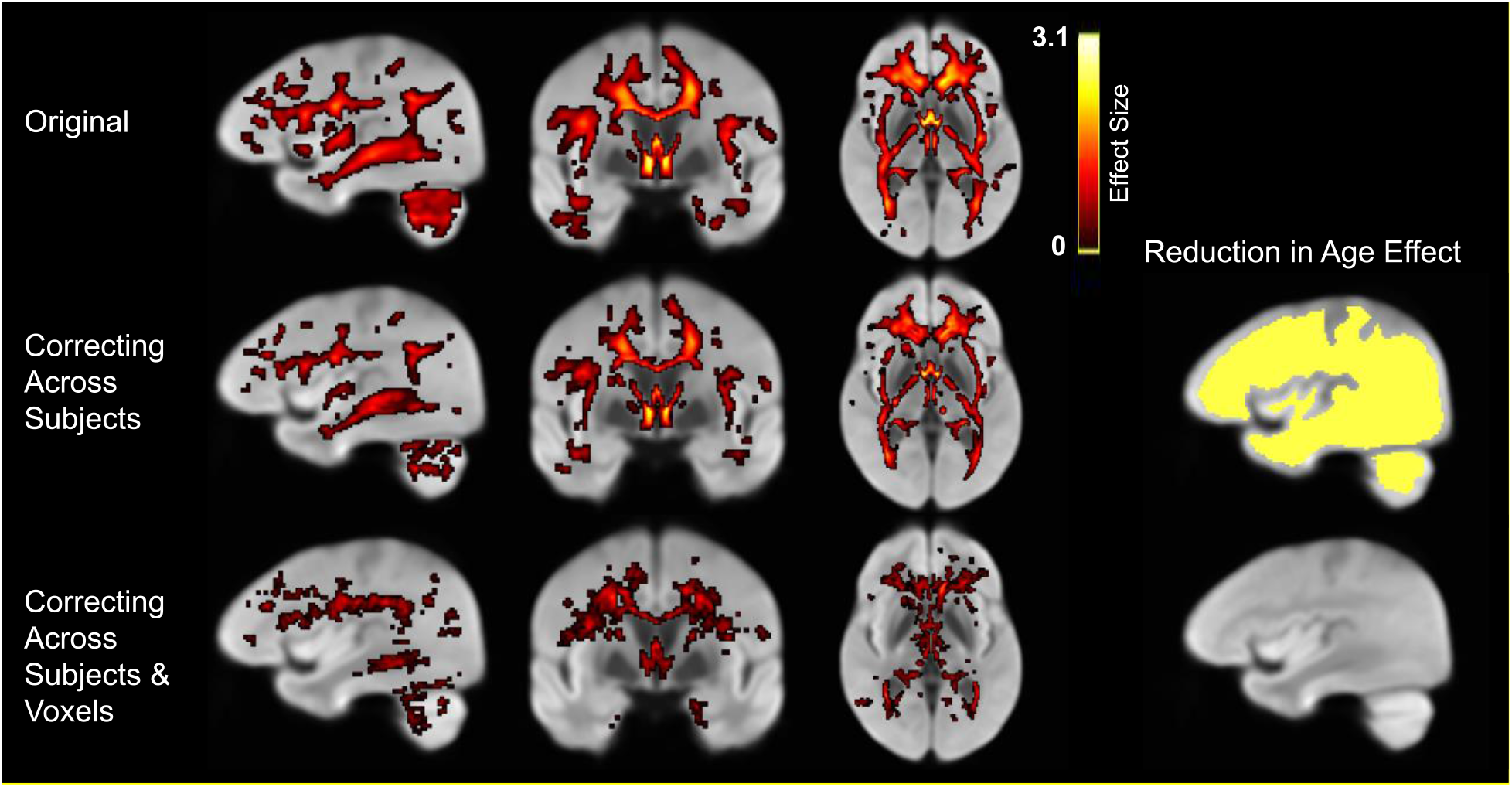
Effects of controlling for complexity. (Left) Apparent age differences in FA in the original data (top), after controlling for CX-FA correlations across subjects (middle), and then across voxels (bottom). (Right) Voxels exhibiting significantly reduced age differences in FA after each stage of correction.

Controlling for within-subject FA-CX correlations across voxels had very little additional effect beyond the effect of controlling for mean CX across subjects (76 additional voxels; 0.02%; Figure 11, right, bottom row), with no discernable spatial pattern across voxels. This correction produced only modest, spatially indiscriminate reductions in apparent age effects (Fig 11, left, bottom row), reducing the number of significant voxels to 45,618 (11.6% of the analysis mask).

This result further suggests that age-associated FA-CX correlations manifest at a relatively global level, with little specificity to particular structures or regions.

### 3.5 Tract Specific DTI analysis controlling for CX

We also evaluated the effects of controlling for CX in comparing tract-wise FA values between age groups (Figure 12 and Table 3). After controlling for CX across subjects, mean FA was significantly lower in older vs. younger adults in bilateral parietal parahippocampal cingulum (*P*=.01 for left and *P*=.020 for right) forceps minor (*P*<.001), fornix (*P*<.001), bilateral IFOF (*P*<.001 for both left and right), bilateral internal capsule (*P*=.003 for left and *P*<.001 for right), bilateral SLF I (*P*<.001 for both left and right), bilateral SLF II (*P*<.001 for left and *P*=.022 for right), and SLF III (*P*=.004 for left and *P*=<.001 for right). In contrast, mean FA was significantly greater in older than younger adults in right subgenual (*P*<.001) and right retrosplenial cingulum (*P*=.002) (not shown).

**Figure 12.**
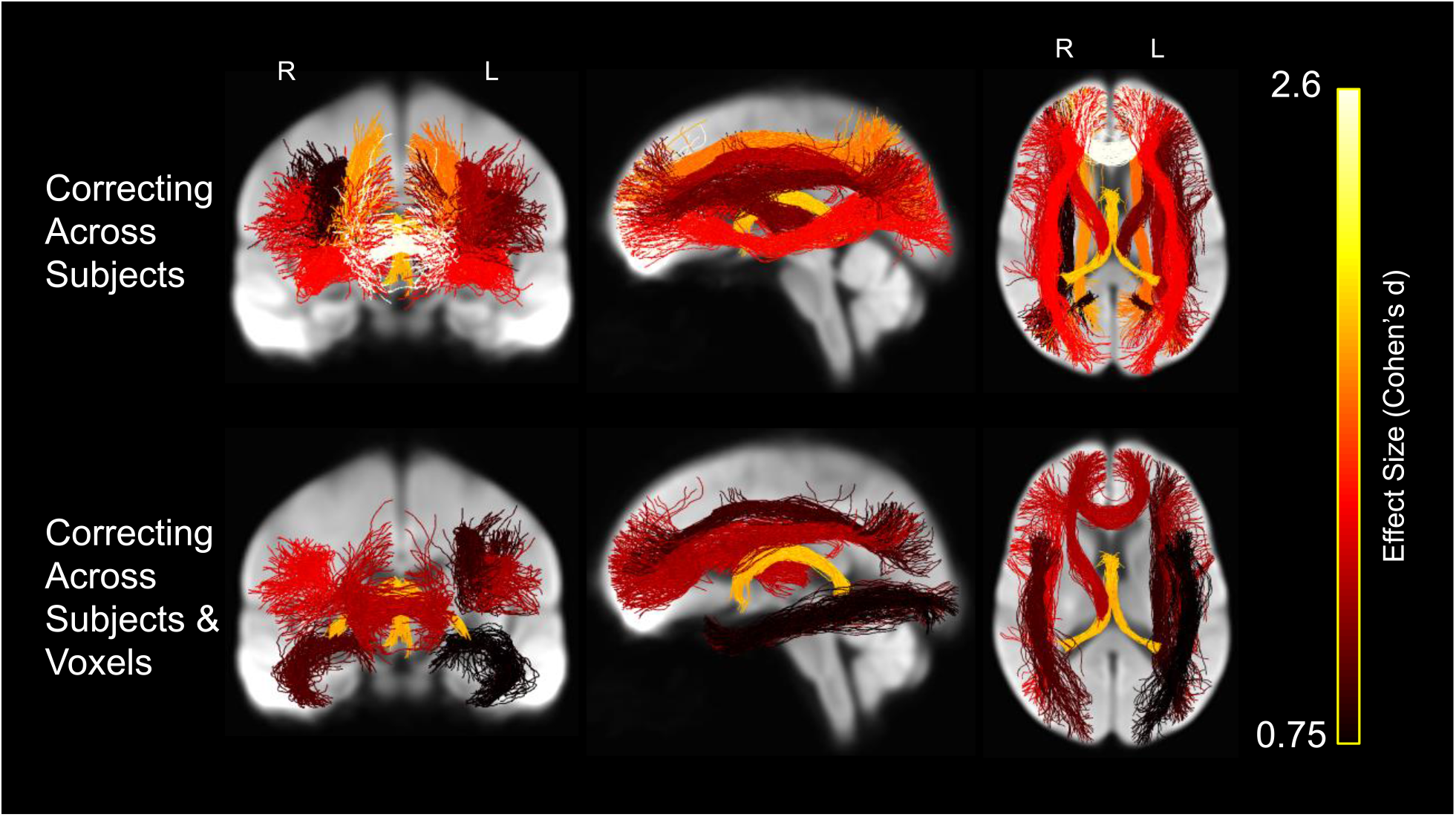
Whole brain projections onto 2-D slices showing white matter tracts in which average fractional anisotropy was significantly lower in the older vs. younger participants after controlling for complexity (FA-CX), displayed on the white matter population template. Streamlines within each tract are colored by the tract’s effect size (Cohen’s d), with brighter colors representing greater effect size.

After further controlling for FA-CX correlations across voxels, FA was significantly lower in older vs. younger adults in forceps minor (*P*<.001), fornix (*P*<.001), bilateral ILF (*P*=.040 for left and *P*=.009 for right), right internal capsule (*P*<.001), left SLF II (*P*=.018), and bilateral SLF III (*P*=<.001 for both left and right). In contrast, mean FA controlling for CX was significantly greater in older than younger adults in right subgenual (*P*=.006) and bilateral retrosplenial cingulum (*P*=.002 for left and *P*=.003 for right) (not shown).

The effects of CX-correction on tract-level FA effects can be appreciated by comparing the first three columns of Table 3. In general, apparent age effects on FA decreased with each stage of correction for FA-CX correlations.

### 3.6 Whole Brain Fixel-Based Analysis

Following the DTI analyses, we evaluated age group differences in the fixel-based analysis metrics: fiber density (FD), fiber cross-section (FC) and combined fiber density and crosssection (FDC) (Figure 13). FD in older adults was significantly lower in the fornix, bilateral anterior internal capsule, forceps minor, body of the corpus callosum and corticospinal tract, relative to younger participants. In addition, older adults exhibited significantly lower FC than younger adults in the cingulum bundle and forceps minor. Last, FDC in older adults was significantly lower in the anterior subcortical white matter regions, including forceps minor, anterior limb of the internal capsule, subgenual cingulum, fornix and IFOF. In general, the FBA-based metrics exhibited a gradient along the anterior-posterior axis with greater age effects in more anterior regions. Furthermore, most of the age effects in anterior brain regions were due to differences in cross-section, rather than density (Figure 13).

**Figure 13.**
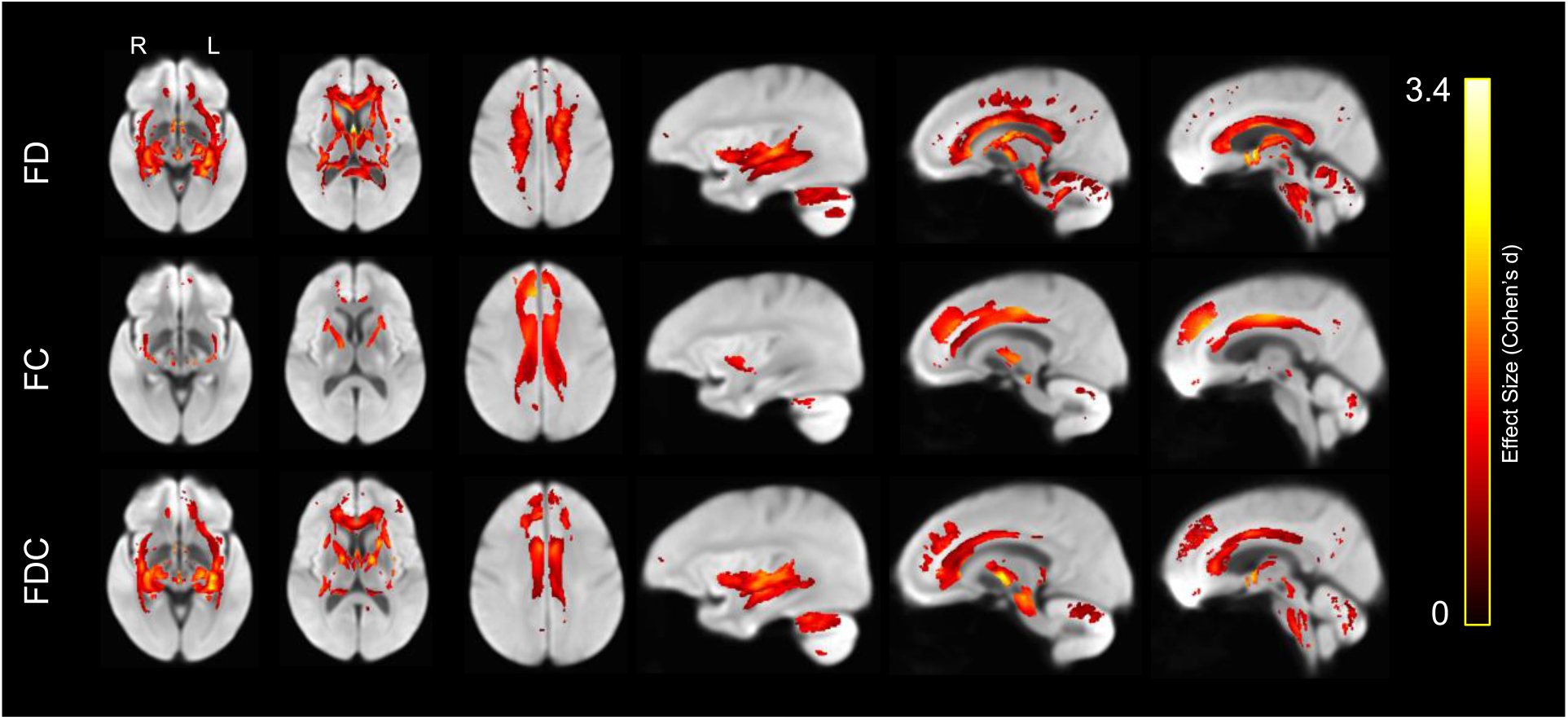
Lower white matter fiber density (FD, top row), fiber cross-section (FC, middle row), and the product of fiber density and cross-section (FDC, bottom row) in older compared to younger adults, displayed on the white matter population template. Colors represent fixels in which the corresponding measure was significantly lower in the older compared with the younger group, with brighter colors representing larger effect size (Cohen’s d).

Importantly, the opposite contrast also revealed a set of white matter regions in which these measures were greater in older participants than in their younger counterparts (Figure 14). Whereas FD in older adults was only greater in bilateral SLF I, FC was greater in forceps major and the body of the corpus callosum. FDC was only greater in forceps major.

**Figure 14.**
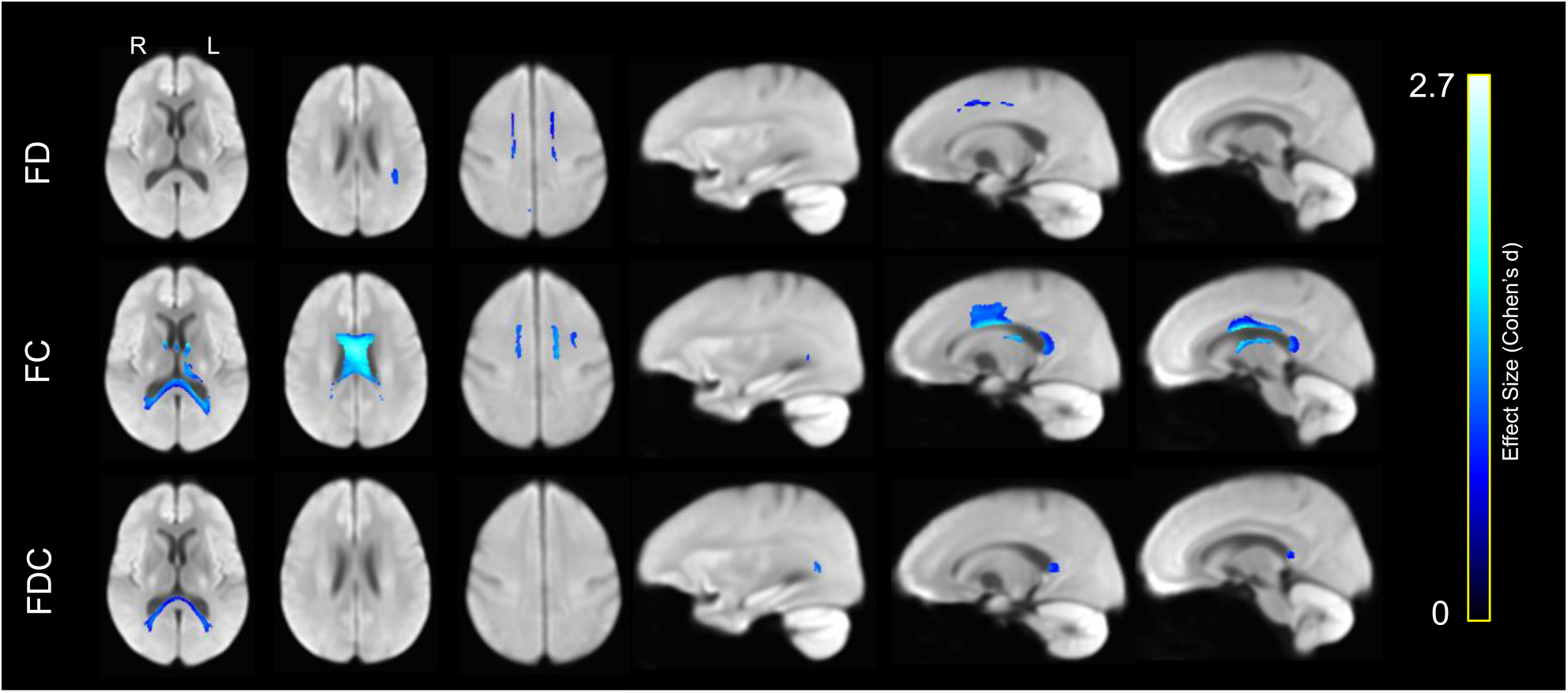
Greater white matter fiber density (FD, top row), fiber cross-section (FC, middle row), and the product of fiber density and cross-section (FDC, bottom row) in older compared to younger adults, displayed on the white matter population template. Colors represent fixels in which the corresponding measure was significantly greater in the older compared with the younger group, with brighter colors representing larger effect size (Cohen’s d).

### 3.7 Tract-Specific Fixel-Based Analysis

Last, we compared mean tract-specific fixel-based parameter values between age groups. As shown in Figure 15 and Table 3, FD was significantly lower in older than younger across multiple tracts: in bilateral temporal (*P*<.001) and right parietal parahippocampal cingulum (*P*=.023), bilateral inferior corticospinal (*P*<.001), forceps minor (*P*<.001), fornix (*P*<.001), bilateral IFOF (*P*<.001), bilateral internal capsule (*P*<.001) and left uncinate fasciculus (*P*<.001). FC was significantly lower in older adults in bilateral retrosplenial (*P*<.001) and subgenual cingulum (*P*<.001), bilateral inferior corticospinal tract (*P*=.003 for left and *P*=.002 for right) and bilateral SLF I (*P*=.005 for left and *P*=.001 for right). FDC was significantly lower in older adults in bilateral retrosplenial (*P*<.001), bilateral subgenual (*P*<.001), bilateral temporal parahippocampal cingulum (*P*<.001), bilateral inferior corticospinal (*P*<.001), forceps minor (*P*<.001), fornix (*P*<.001), right IFOF (*P*=.009), right ILF (*P*=.030), bilateral internal capsule (*P*<.001) and bilateral uncinate fasciculus (*P*<.001 for left and *P*=.030 for right). FD was greater in older adults in left SLF I (*P*=.013) and FC was greater in older adults in fornix (*P*<.001) (not shown).

**Figure 15.**
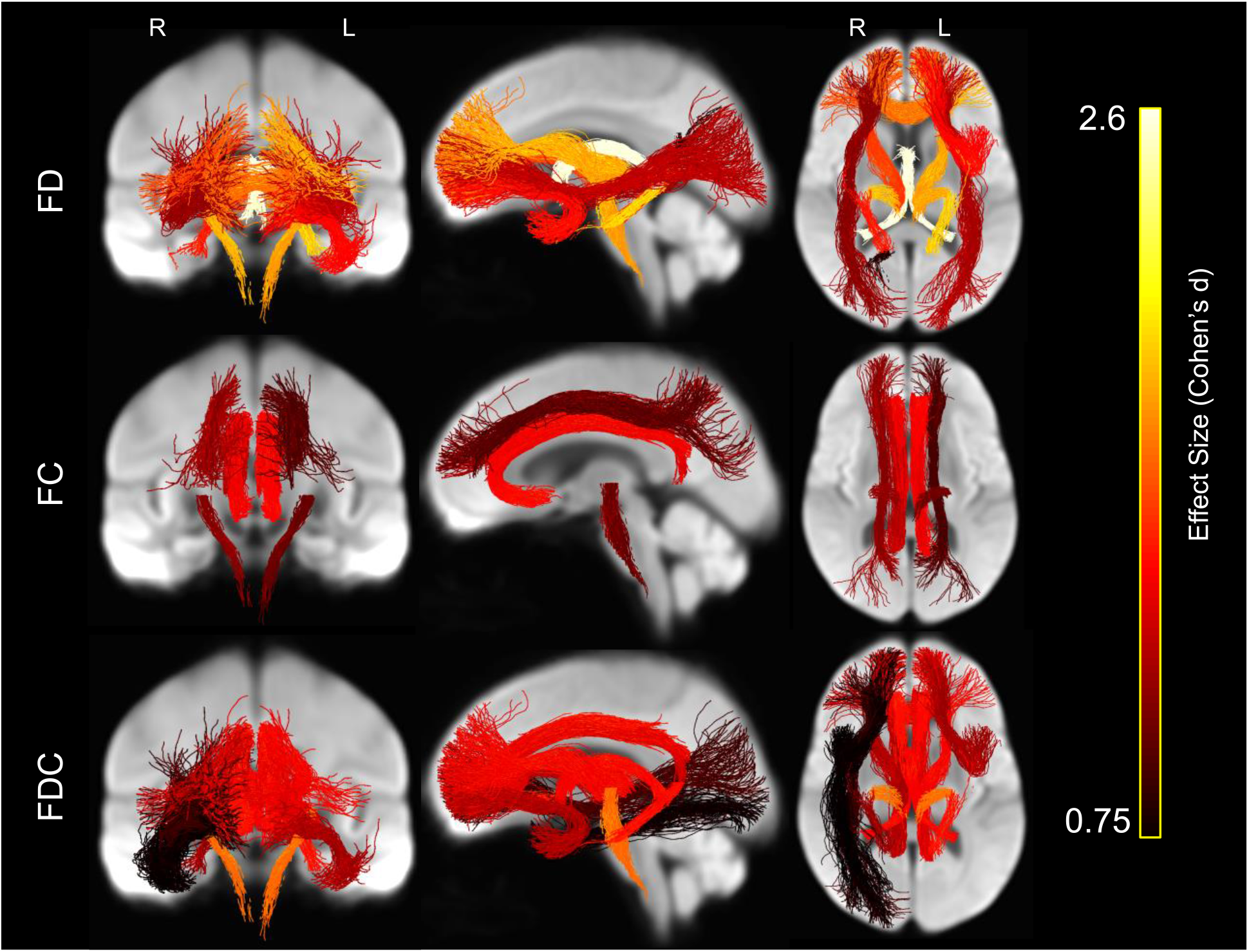
Whole brain projections onto 2-D slices showing white matter tracts in which average fiber density (FD, top row), average fiber cross-section (FC, middle row), and the product of average fiber density and cross-section (FDC, bottom row) was significantly lower in the older vs. younger participants, displayed on the white matter population template. Brighter colors represent greater effect size (Cohen’s d).

### 3.8 Summary of Age Effects

Table 3 presents a summary of the effects of age on all the measures analyzed here: FA, FA after controlling for complexity, FD, FC, and FDC. The table presents significant results from analyses in which age was treated as a dichotomous variable. We repeated these analyses while treating age as a continuous variable and all 84 significant results replicated.

## 4. Discussion

The present study establishes multiple notable effects regarding age-related differences in white matter structure and organization. First, by combining traditional DTI modeling of dMRI data with newer fixel-based approaches we show that some, but not all age differences in FA, are influenced by local multi-fiber geometry (crossing fibers) within individual voxels. Indeed, age-based effects of fiber complexity on FA manifest rather globally, leading to overestimates of age effects on FA throughout the brain’s white matter. Second, by separately accounting for differences in fiber density and cross-section, the results from fixel-based analyses afford new insight into the micro- and macro-structural nature of age differences in white matter pathways beyond what has been reported in prior DTI aging studies. Specifically, age differences in fiber density were prominent in fornix, bilateral anterior internal capsule, forceps minor, body of the corpus callosum and corticospinal tract, while age differences in fiber cross section were largest in cingulum bundle and forceps minor. Third, we report DTI and fixel-level findings that reveal a more heterogeneous pattern of age differences than are commonly reported, independent of crossing fibers.

### 4.1 Modeling crossing fibers changes age differences in FA

This is one of the few extant reports combining common tensor approaches with newer fixel-based analysis and the only such report examining aging. Crucially, FA and CX, an index of intravoxel multi-fiber complexity, exhibited a significant inverse relationship (r = −0.81) in voxels exhibiting a significant age effect on FA. Following initial results showing widespread age differences in FA, we subsequently re-evaluated these effects on FA while statistically controlling for CX (Riffert et al. 2014). The original more expansive pattern of voxels with significantly lower FA in older adults was considerably attenuated after controlling for CX. FA in forceps minor, fornix, and anterior limb of the internal capsule remained significantly lower in older adults, but the age differences in multiple association fiber tracts (i.e., parts of the IFOF, ILF, SLF, cingulum, and uncinate) were rendered nonsignificant after controlling for CX. All or major portions of each of these association tracts also did not exhibit significant effects for fixel-based metrics, supporting the notion that some of the observed age differences in FA are due to differences in multi-fiber organization (i.e., the relative density of crossing fibers). The centrum semiovale, which includes the intersection of the SLF, CST, and corpus callosum, is one region in which tensor model parameters are already known to be strongly confounded by crossing fibers (Pierpaoli and Basser 1996; Rokem et al. 2015), but our results suggest that numerous other areas are affected as well.

Interestingly, we found that FA-CX correlations manifested both across subjects (i.e., correlations between mean CX and FA) and within subjects (i.e., correlations across voxels). These results suggest that apparent age effects on FA are confounded with age-related differences in CX, resulting in an over-estimation of apparent FA differences throughout the brain. Importantly, CX-FA correlations primarily influenced apparent age effects at the level of individual differences in mean FA, suggesting that apparent age differences in FA partially reflect relatively global differences in CX. While sensitive to age differences, the observed lack of specificity in FA underscores long held concerns about interpreting this parameter as a biologically meaningful measure of white matter ‘integrity.’ One possibility is that both observed age-associated differences in CX and FA likely reflect more variable, higher dimensional changes in fiber density and cross-section across multiple underlying fiber populations within a voxel (Douaud et al. 2011; Yang et al. 2013). However, in white matter voxels with crossing fibers one cannot know whether decreases in FA reflect reduced signal from the largest fiber population, increased signal from orthogonal populations, or other combinations of intra-voxel influences (Riffert et al. 2014). Further work is needed to better understand how older age may differentially affect fiber density and cross-section in secondary and tertiary crossing fibers, relative to primary fibers in a voxel.

### 4.2 Integrating Tensor and Fixel Results

The FA analyses revealed age differences across numerous white matter regions, even after correcting for crossing fibers. The fixel-based analyses afford new insight into the orientationally specific, micro- and macro-structural white matter characteristics reflected by these FA differences. For example, we found that younger adults exhibited significantly higher FA than older participants in forceps minor and fornix, independent of crossing fibers. However, the fixel-based analysis showed that these effects may reflect different underlying age differences in white matter organization. Whereas younger adults had significantly higher fiber density in both these regions, the measure of fiber cross section in the fornix (but not forceps minor) was greater for older adults. One interpretation is that the fornix shows signs of atrophy and loss of packing density even in normal aging (Peters et al. 2010). Younger adults also had significantly higher FD than older adults in other regions, including right parietal parahippocampal cingulum, bilateral anterior IFOF, and bilateral internal capsule. These findings suggest that the original FA results were likely due to age differences in both multi-fiber organization and fiber density. Fiber cross-section in SLF I was higher in younger adults, suggesting that the original FA results may have been due to age differences in both multi-fiber organization and macroscopic measures of fiber cross-section.

The fixel-based analysis also uncovered age differences that were not observed in the DTI analysis. In particular, fiber cross-section and the product of fiber density and cross-section were both significantly lower in the older group in retrosplenial and subgenual aspects of the cingulum bundle, but FA and fiber density were not. We also observed greater fiber cross-section, but lower fiber density, in the body of the corpus callosum, forceps major, and fornix in older participants compared to younger adults. These results are consistent with histological findings showing a preferential loss of smaller diameter axonal fibers in older age (Aboitiz et al. 1992; Bowley et al. 2010; Marner et al. 2003; Peters et al. 2010).

We also found that the FBA-based metrics exhibited a gradient along the anterior-posterior axis with greater age effects in more anterior regions. Furthermore, most of the age effects in anterior brain regions were due to differences in cross-section, rather than density (Figure 13). One potential interpretation of this result is that as we age, the number of fibers in white matter tracts in the frontal lobe and anterior limbic regions declines, but that the density of the fibers in those bundles does not.

### 4.3 Heterogeneous age differences

The present results revealed several regions in which older adults had higher FA, FC, and FDC than younger participants, highlighting the multidimensional nature of white matter aging. Like some other prior reports, we observed greater FA in the superior cerebellar peduncles, as well as in the external capsule and in the cingulum bundle of older adults (Kanaan et al. 2016). Our results are also consistent with reports of longitudinal increases in FA in early-developing white matter regions in middle-aged and older adults (Bender and Raz; 2015; Bender et al. 2016). We also found greater FC and FDC in the splenium of the corpus callosum/forceps major in older adults compared to younger adults, but no age differences in fiber density. Greater FC and FDC may reflect fiber populations with greater diameters, either in the size of the axon bundle or degree of myelination. The visual system, of which the forceps major is a part, develops early in life and has high levels of use and automaticity. Our findings of higher FC and FDC in the forceps major are consistent with the hypothesis that earlier-developing white matter pathways supporting consistently utilized and automated behavior (like forceps major of the visual pathway) may be less vulnerable to age-related declines (Bender et al. 2016; Karolis et al. 2019).

### 4.4 Limitations and future directions

One obvious limitation of the current study is that the data are cross-sectional rather than longitudinal. Some of the observed differences could therefore be due to cohort effects rather than within-person change as a result of age. Also, our sample of older adults were better educated than most and may not be representative of a less educated population. Similarly, we did not collect information on health and physiological function in these participants; future studies could evaluate how vascular and metabolic factors influence these newer white matter parameters in older adults.

We also performed our analysis on single-shell b = 1000 s/mm^2^ dMRI data, rather than data with multiple or higher b-values, which would have improved angular resolution and biological interpretation of underlying observed effects. When diffusion sensitivity is high (b > 2000 s/mm^2^), the integral over an fODF lobe (“apparent fiber density”) is proportional to fiber density within an associated fiber population (Raffelt et al. 2012). With less diffusion sensitization, as used in this study (b = 1000 s/mm^2^), this metric is also sensitive to extra-axonal diffusion, and therefore may also reflect axonal hindrance of extra-cellular diffusion and other microstructural factors. Nevertheless, our results, as well as those of Toselli et al. (2017), demonstrate that fixel-based approaches have advantages over DTI methods when analyzing single shell data like those collected here. Furthermore, our ability to observe the results reported here using data collected in 32 gradient directions demonstrates that the results are quite robust and suggest the promise of applying this method to existing single-shell lower b-value data. Of course, it would be preferable to collect higher resolution multi-shell data in the future (as we are now doing).

We also note that the “complexity” (CX) metric used in this study does not fully capture all geometric factors (e.g., fiber angle) that contribute to FA or the apparent “complexity” of an fODF. Nevertheless, the strong correlations between FA and CX observed in this study demonstrate the need to control for crossing fiber influences whenever investigating group differences in tensor-based parameters.

Finally, it is important to note that our findings build on prior studies using non-tensor dMRI approaches. For example, Tuch (Tuch 2004) developed a method (termed q-ball imaging) that was able to distinguish multiple differentially oriented fiber bundles in individual voxels. Assaf and Baker (Assaf and Baker 2005) (also see De Santis et al. 2014) developed a composite hindered and restricted model of diffusion (CHARMED) data that produced estimates of fiber orientation that had less angular uncertainty than the traditional diffusion tensor model, and De Santis et al. (De Santis et al. 2012) used this model to investigate the biophysical correlates of diffusional kurtosis imaging (DKI). More recently, Toschi et al. (Toschi et al. 2020) used the CHARMED model to examine age-related changes in white matter and found that these changes begin earlier in men than in women and they affect more frontal regions. Billiet et al. (Billiet et al. 2015) collected multishell diffusion MRI data as well as multiexponential T2 relaxation (MET2) data in a sample of 59 adults ranging in age from 17 to 70 and reported evidence that age-related frontal decrease in FA may reflect increased axonal dispersion and not demyelination.

In conclusion, DTI has been a useful tool for investigating white matter differences in age groups. However, its main metric FA can be difficult to interpret as it is sensitive to multiple factors. Here we used complexity to account for multi-fiber organizational differences and fixel-based analysis to investigate the intricacies underlying age differences in FA. Like other studies that have used multiple methods to investigate age-related changes in white matter, the current study demonstrates the power of employing different approaches to white matter analysis alongside DTI in order to gain insight into white matter differences associated with aging.

## Acknowledgements

This work was supported by a grant from the National Institutes of Health to TAP (R01AG050523).

