## Supplemental Figure 1 for "Age-related differences in white matter: Understanding tensor-based results using fixel-based analysis"

### Supplementary Figures

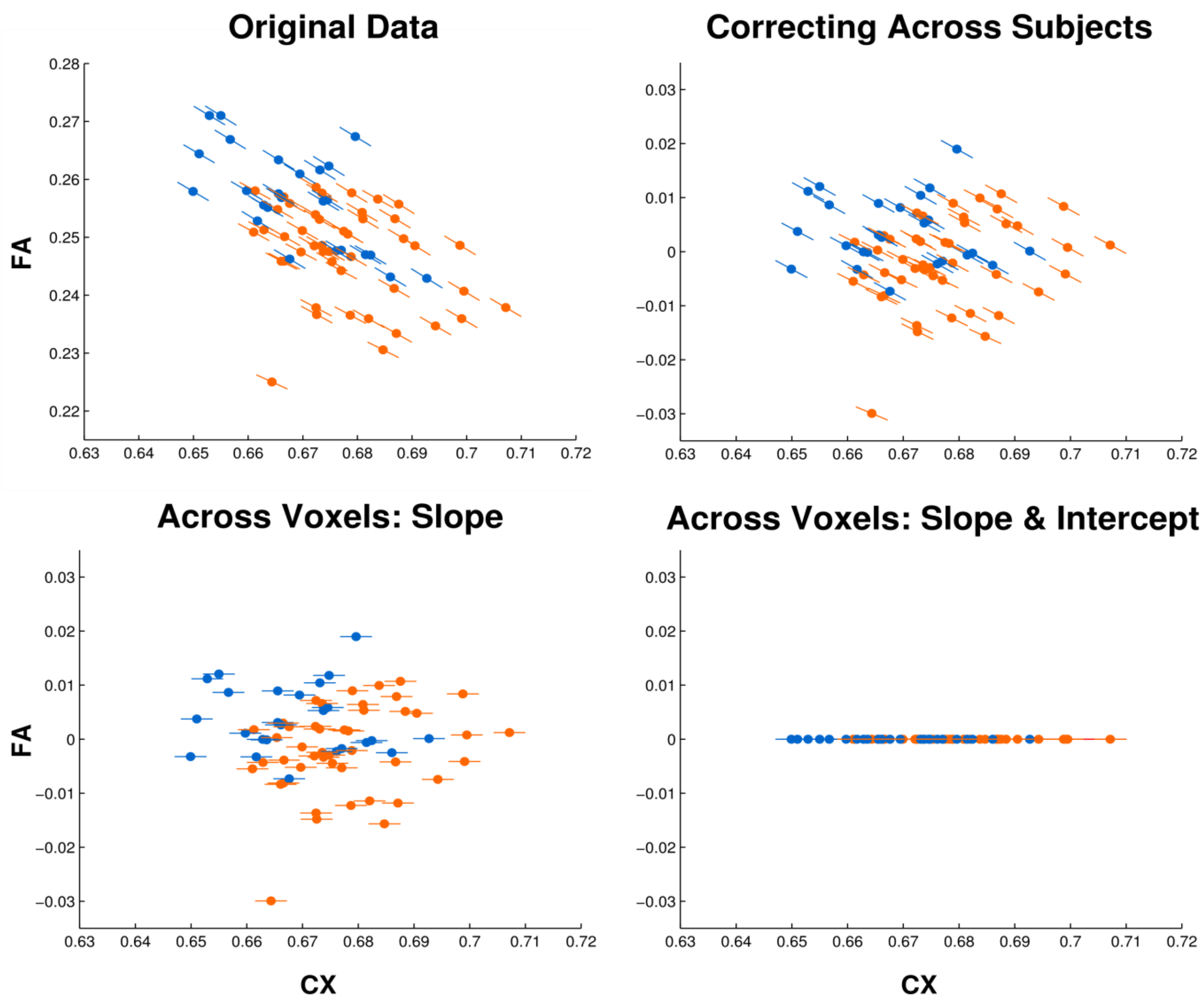


**Figure S1.** Corrections for CX-FA correlations. The upper left panel shows the correlational structure of the original data, with points indicating mean values for each subject within the white matter mask, and lines indicating the slopes of voxel-wise regressions within each subject. As shown in the upper right panel, correcting for CX-FA correlations at the subject level shifts mean FA values towards 0, but does not completely abolish differences in mean FA between the young (blue) and old (orange) groups. Subtracting the slope term of each voxel-wise regression eliminated the CX-FA dependence within each subject (lower left) without removing residual individual differences in mean FA. As shown in the lower right panel, subtracting the intercept term would abolish these residual differences in mean FA.

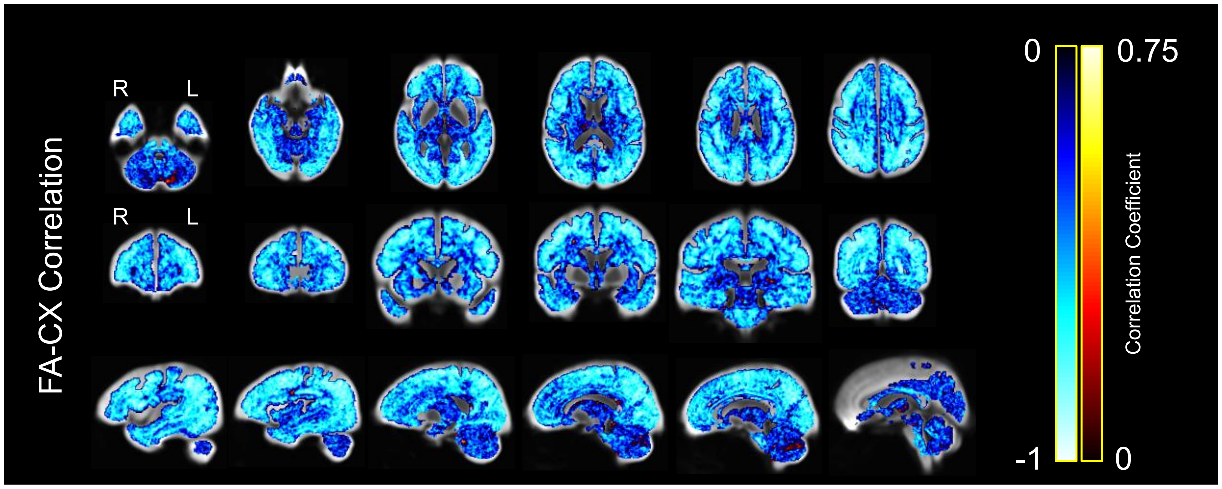


**Figure S2.** Average correlation between FA and CX across subjects within each voxel of total white matter for young subjects only, displayed on the white matter population template. Colors represent the strength of the negative or positive correlation coefficient for each voxel, with brighter colors representing stronger correlation coefficients.

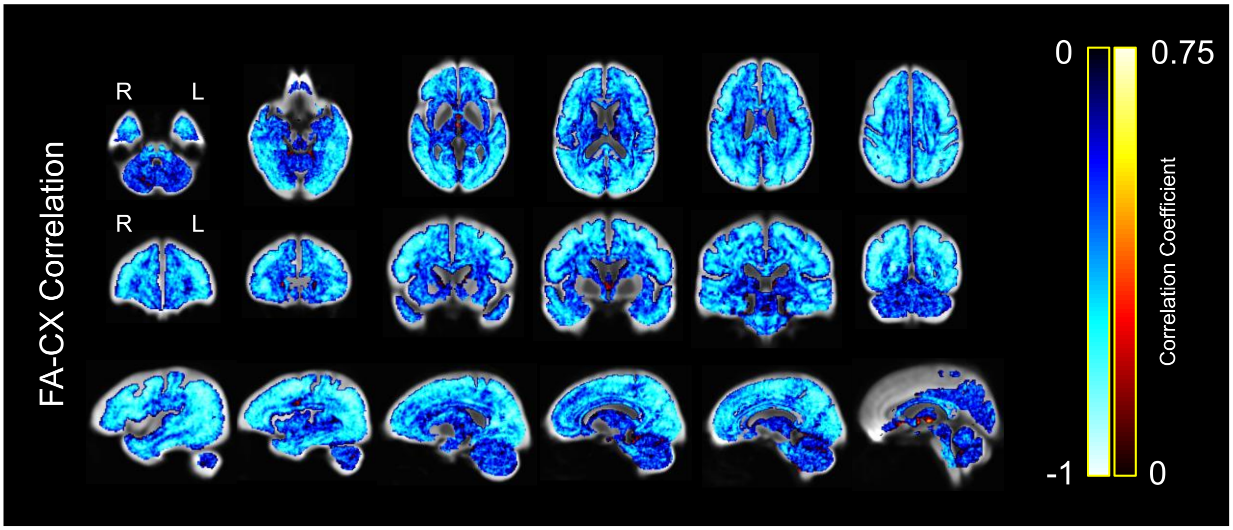


**Figure S3.** Average correlation between FA and CX across subjects within each voxel of total white matter for old subjects only, displayed on the white matter population template. Colors represent the strength of the negative or positive correlation coefficient for each voxel, with brighter colors representing stronger correlation coefficients.

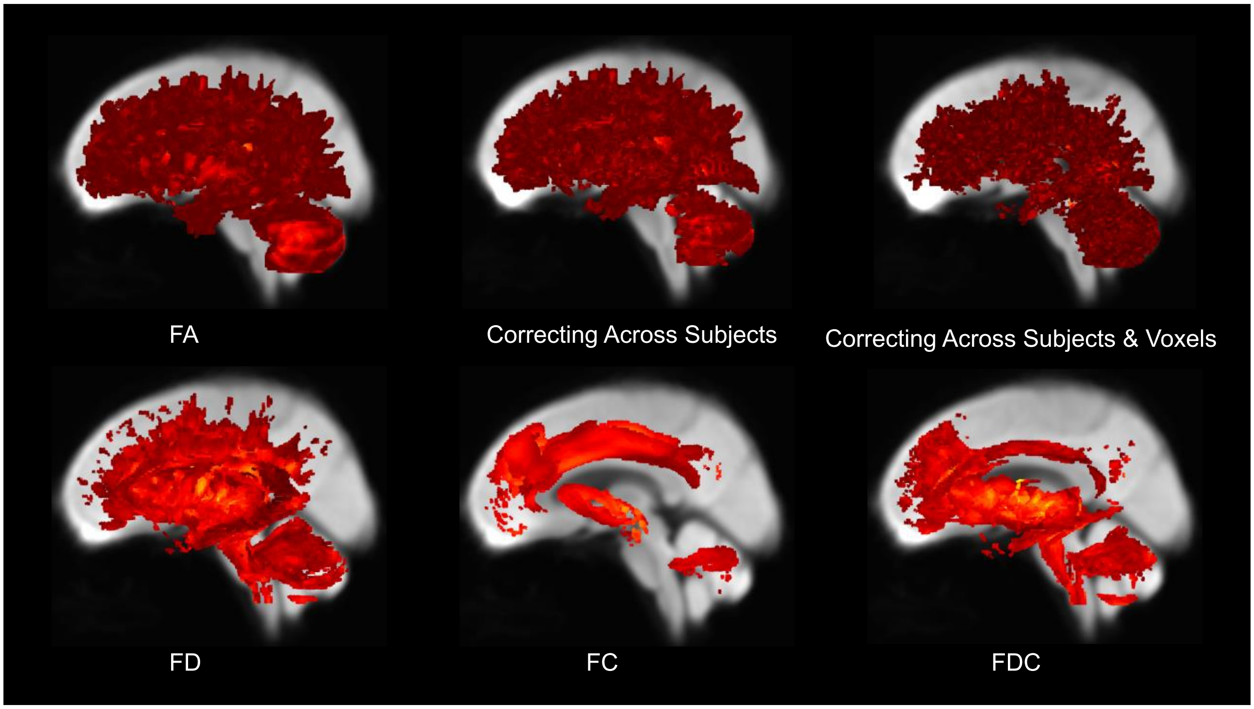


**Figure S4.** Lower white matter fractional anisotropy (FA), FA after controlling for complexity (CX), fiber density (FD), fiber cross-section (FC) and fiber density and cross-section (FDC) in older compared to younger adults, displayed on the white matter population template. Streamline segments correspond to voxels or fixels in which FA, FA-CX, FD, FC, or FDC respectively were significantly lower in the older compared with the younger group.
